# Cell-to-cell variation in defective virus expression and effects on host responses during influenza virus infection

**DOI:** 10.1101/487785

**Authors:** Chang Wang, Christian V. Forst, Tsui-wen Chou, Adam Geber, Minghui Wang, Wissam Hamou, Melissa Smith, Robert Sebra, Bin Zhang, Bin Zhou, Elodie Ghedin

## Abstract

Virus and host factors contribute to cell-to-cell variation in viral infections and determine the outcome of the overall infection. However, the extent of the variability at the single cell level and how it impacts virus-host interactions at a systems level are not well understood. To characterize the dynamics of viral transcription and host responses, we used single-cell RNA sequencing to quantify at multiple time points the host and viral transcriptomes of human A549 cells and primary bronchial epithelial cells infected with influenza A virus. We observed substantial variability of viral transcription between cells, including the accumulation of defective viral genomes (DVGs) that impact viral replication. We show a correlation between DVGs and viral-induced variation of the host transcriptional program and an association between differential induction of innate immune response genes and attenuated viral transcription in subpopulations of cells. These observations at the single cell level improve our understanding of the complex virus-host interplay during influenza infection.

**IMPORTANCE:** Defective influenza virus particles generated during viral replication carry incomplete viral genomes and can interfere with the replication of competent viruses. These defective genomes are thought to modulate disease severity and pathogenicity of the influenza infection. Different defective viral genomes also introduce another source of variation across a heterogeneous cell population. Evaluating the impact of defective virus genomes on host cell responses cannot be fully resolved at the population level, requiring single cell transcriptional profiling. Here we characterized virus and host transcriptomes in individual influenza-infected cells, including that of defective viruses that arise during influenza A virus infection. We established an association between defective virus transcription and host responses and validated interfering and immunostimulatory functions of identified dominant defective virus genome species *in vitro*. This study demonstrates the intricate effects of defective viral genomes on host transcriptional responses and highlights the importance of capturing host-virus interactions at the single-cell level.

## INTRODUCTION

The productivity of viral replication at the cell population level is determined by cell-to-cell variation in viral infection [1]. The genetically diverse nature of RNA viruses and the heterogeneity of host cell states contribute to this inter-cell variability, which can impact therapeutic applications [2, 3]. Although previous studies of cell-to-cell variation during viral infection have mainly centered on non-segmented viruses such as poliovirus [2, 4], vesicular stomatitis virus (VSV) [5, 6], or dengue (DENV) and zika (ZIKV) viruses [7], heterogeneity across cells may be further complicated for viruses with a segmented genome, such as influenza A virus (IAV). IAV-infected cells display substantial cell-to-cell variation as it pertains to relative abundance of different viral genome segments [1, 8], transcripts [9], and their encoded proteins [10], which can result in non-productive infection in a large fraction of cells.

Defective interfering (DI) particles readily generated during successive high multiplicity of infection (MOI) in cell culture passages [11–14] and observed in natural infections [15, 16] are likely by-products of the IAV replication process and have a significant effect on productive infection. DI viruses have incomplete viral genomes with large internal deletions and they possess the ability to interfere with the replication of infectious viruses. Because DIs diminish the productivity of infectious progenies and could impact disease outcome, they are of great interest for therapeutic and prophylactic purposes (reviewed in [17]). A specific influenza DI (i.e., DI244) was shown to effectively provide prophylactic and therapeutic protection against IAV infection in mice [18]. One of the ways influenza DIs are thought to modulate viral infections is through interaction with a cytosolic pathogen-recognition receptor (PRR), RIG-I, essential for interferon (IFN) induction [19]. Besides enhanced IFN induction during infection with DI-rich influenza virus populations observed *in vitro* and *in vivo*, it has been assumed that DI virus could also compete with standard virus for cellular resources (reviewed in [17, 20]).

While diverse DI viruses can arise during IAV infection [21], the emergence and accumulation of distinct DIs, as well as other defective virus genomes (DVGs), has not been characterized at a single cell resolution, although the diversity of DIs present could be contributing to the observed cell-to-cell variation in host transcription.

We probed viral and host transcriptomes simultaneously in the same cells using single-cell RNA-seq to monitor host-virus interactions in cultured cells over the course of the infection. This data established a temporal association between the level of viral transcription and the alteration of the host transcriptome, and characterized the diversity and accumulation of DVG transcripts.

## RESULTS

### Cell-to-cell variation in virus gene expression

To determine how both the viral and host cell transcriptional programs relate to each other over the course of an influenza infection, we infected two cell types—the adenocarcinomic human alveolar basal epithelial A549 cell line and human primary bronchial epithelial cells HBEpC—at high multiplicity of Infection (MOI; 5) with A/Puerto Rico/8/34(H1N1) (PR8) and performed single cell and bulk RNA-seq expression analyses. A high MOI infection ensures that virtually all the cells can be rapidly infected, promotes the accumulation of DVGs, and consequently enables the characterization of both host response and DVG diversity. We first determined the percentage of reads that uniquely aligned to viral genes from the total number of mapped reads to obtain the relative abundance of virus transcripts within cells at each time point. Similar to what has been observed at early stages of infection during a low MOI infection of IAV [9], the relative abundance of virus transcripts was heterogeneous across cells from both cell types, with 0 to 70% of the total reads in each cell being derived from virus transcripts and the relative abundance of these transcripts increasing over time (Supplementary Fig. 1a). The same trend was also seen when analyzing segment-specific virus transcripts within individual cells over the course of the infection (Supplementary Fig. 1b).

### Heterogeneity of defective virus segment expression across the cell population

Since DVGs are known to accumulate in cell culture and can serve as templates for transcription, we characterized their abundance and diversity by examining gap-spanning reads in the sequencing data. To quantify the relative abundance of DVGs identified, we calculated the frequency of each unique DVG transcript type (determined by 5’ / 3’ gap coordinates) across single cells and the ratio of these to non-gap-spanning transcripts derived from the corresponding viral segments (i.e., DVG/FL ratio). Although DVG transcripts could be detected in all viral segments, those originating from the polymerase segments (i.e., PB2, PB1, and PA) occurred with the highest abundance and frequency in both cell types over the course of the infection (Supplementary Fig. 2), consistent with observations made in other studies [15, 22]. We thus focused our analyses on the detection of DVG transcripts derived from those segments. We collected reads with large internal deletions spanning ≥1000 nucleotides (nt) (Fig. 1a) and identified the junction coordinates of these gap-spanning reads. We observed a diverse pool of DVG transcripts, including some shared with the viral segments (vRNA) from the PR8 stock (Supplementary Fig. 3-5). The sizes for the majority of these DVG transcripts are estimated to be between 300nt and 1000nt. To evaluate the accumulation of DVG transcripts over the course of the infection, we compared changes in the ratio of DVG transcripts to full-length (FL) transcripts derived from a given polymerase segment (DVG/FL) across time points. The DVG/FL ratios increased significantly between the early (6hpi) and late (24hpi) stages of the infection (p < 2.2 x 10^-16^) and displayed a high level of heterogeneity across single cells from both cell types (Supplementary Fig. 6).

**Figure 1.**
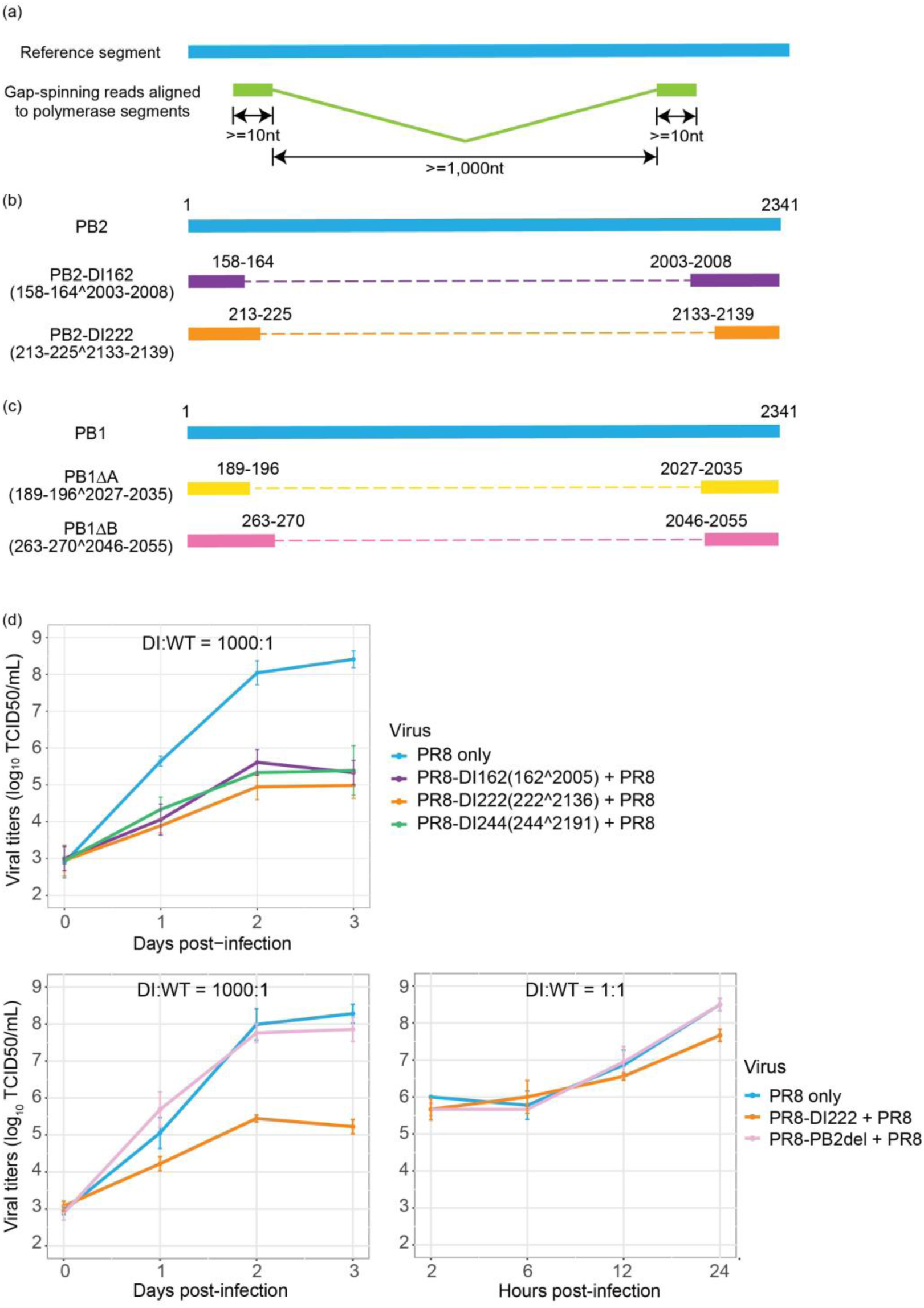
Characterization of DVG transcripts and validation of interfering capacity. **(a)** Selection criteria for reads considered to be representatives of DVGs. Given the length of the reads, we required that each portion of the gap-spanning reads were at least 10 bp in length. Considering that DVGs typically lose approximately 80% of their original sequences and considering the distribution in the size of the deletions detected in this study, we also required that the size of the gap should be at least 1 kb. **(b-c)** Schematic diagram showing two types of defective viral transcripts derived from **(b)** PB2 and **(c)** PB1 segments, respectively. The caret “^” denotes a deletion event and the numbers before and following it indicate the coordinates of the junction sites. **(d)** Interfering capacity of generated defective interfering particles (DIs) carrying identified deletion sites in the PB2 segment during co-infection with the WT-PR8 virus. A549 cells were co-infected with WT virus and one type of DI virus at a ratio of 1:1000 or 1:1. Viral titers were measured by the TCID50 assay on MDCK cells. Four types of defective viruses were tested, including two DI viruses identified in this study, a previously reported DI244 virus [18] that has been considered as a potential candidate for antiviral therapy [17], and a PB2 polymerase defective PR8 virus mutant (i.e., PB2del) that carries 1 nucleotide deletion in the coding sequence of the PB2 gene. For each type of DI virus, the information about the junction sites was denoted in parentheses. The error bar representing the standard deviation of the mean was obtained from three biological replicates. Each panel shows an independent interference assay.

Interestingly, the junction sites aggregated in specific genomic regions, as they systematically occurred in a high percentage of cells. We identified two predominant types of DVG transcripts corresponding to PB2 and PB1 defective segments (Fig. 1b-c and Supplementary Fig. 7-8). These same DVG transcripts were present in the stock, increased in prevalence over the course of the infection, and were conserved in different cell types and at different MOIs (Supplementary Fig. 9-10), indicating likely carryover from the stock virus rather than *de novo* formation in each cell type. For PA, one particular DVG type that was present in the stock was found in A549 and MDCK cells at different MOIs, but not in HBEpC cells. Its prevalence was also much lower than for PB2 and PB1 DVGs (Supplementary Fig. 11-12). The dominant DVG PB2 and PB1 transcripts derived from corresponding defective segments in both the stock and the infected cells showed stable relative abundance across individual cells over the course of the infection, suggesting the persistence of the defective viral segments.

To determine if any of the inferred defective polymerase segments could lead to the generation of defective interfering particles (DIs), we tested their interference potential experimentally at the bulk level. We generated clonal PB8-DI162 and PR8-DI222 viruses carrying the deletion sites in the PB2 segment identified as those in two dominant PB2-DVG species, and then co-infected each of the DI viruses with PR8 wild-type (WT) virus in A549 cells. Both PR8-DI162 and PR8-DI222 viruses inhibited the productivity of WT virus as well as PR8-DI244, known to be an effective DI [18] (Fig. 1d). To confirm that inhibition is specific to the DIs and not due to competition between co-infecting strains, we also tested a full-length PR8 virus with a frame-shifting point mutation in the PB2 gene. Co-infection with this mutant virus did not affect the productivity of WT virus (Fig. 1d). The inhibitory effect of PR8-DI222 virus could still be detected at a DI/WT ratio of 1:1 (Fig. 1d). This confirmed the interfering ability of the two types of defective PB2 segments identified in the single cell data. The interference test was not intended to mimic a single-cell scenario given the difficulties in recreating single-cell infection conditions at a bulk level.

### Cell-to-cell variation in concurrent virus-host transcriptional changes over the course of the infection

Viral infection can trigger massive changes in the host transcriptional program, including the activation of the interferon (IFN) response. Since the induction of IFNs is also subject to cell-to-cell variation [23], we first evaluated the expression frequency of type I (i.e., IFN beta) and III (i.e., IFN lambda) IFNs to determine the extent of the variation during infection. While these were significantly differentially expressed over the course of the infection at the population level, as measured by bulk RNA-seq (Fig. 2a), the expression of IFNs was only detected in less than 3% of cells until the late stage of infection, although this proportion increased over time (Fig. 2b and Fig. 2c). The same expression profile was observed for a number of IFN-stimulated genes (ISGs), such as *RSAD2*, *CXCL10*, *GBP4*, *GBP5*, *IDO1*, and *CH25H* in A549 cells, and *IFI44L*, *CMPK2*, *IFIT1*, *BST2*, *OASL*, and *XAF1* in HBEpC (Fig. 2). In silico pooling of the single cell data to mimic the population level measurement resembles the bulk RNA-seq data, thus excluding the possibility of substantial technical limitation of single-cell RNA-seq (Fig. 2a).

**Figure 2.**
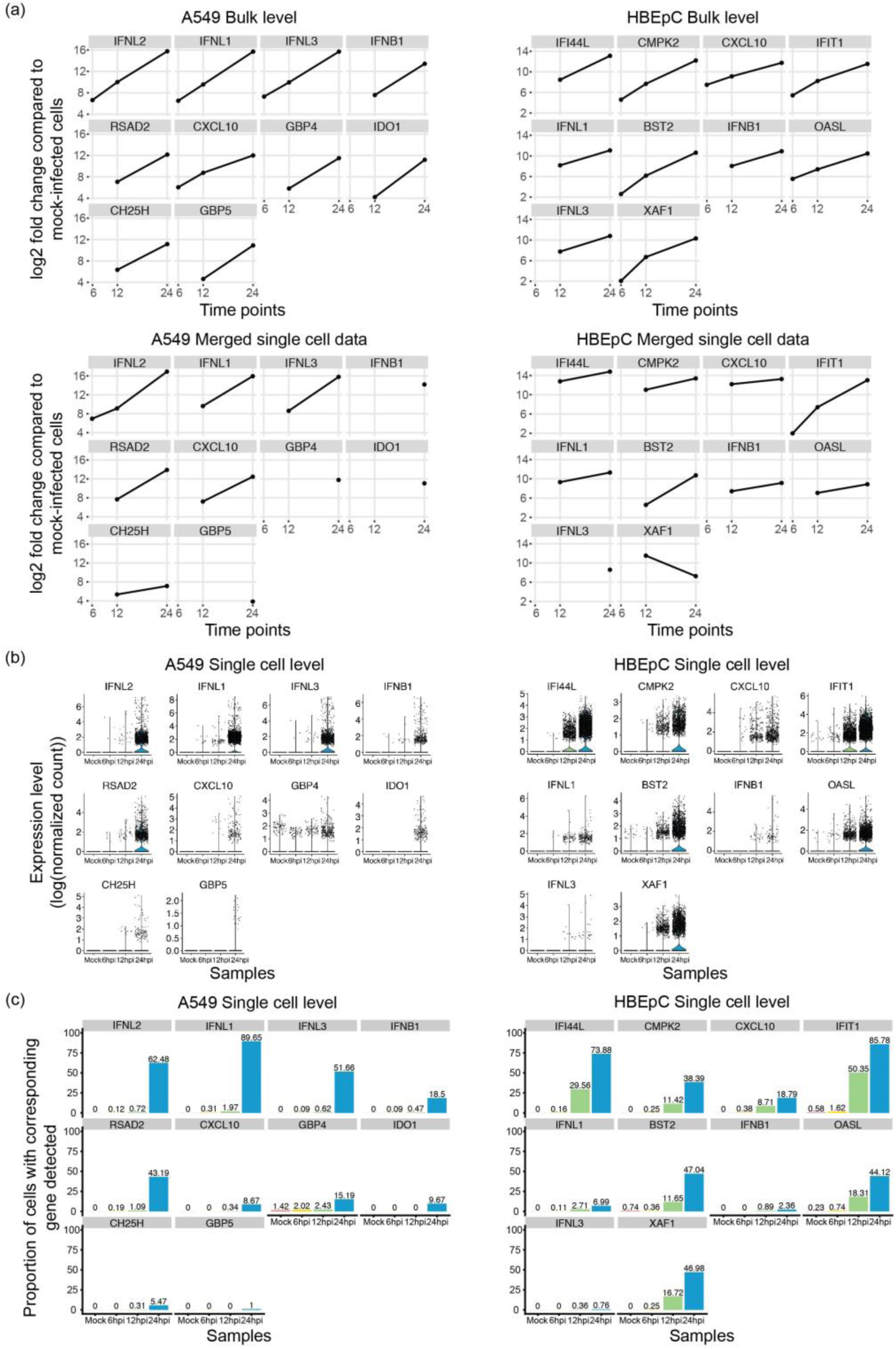
Comparison of the induction of interferons (IFNs) and interferon-stimulated genes (ISGs) at bulk and single-cell levels. **(a)** Log2 fold changes of the top 10 differentially expressed genes with the greatest fold change by 24hpi, including type I and III IFNs and some ISGs, compared to mock-infected cells detected in A549 and HBEpC cells over the course of the infection at the bulk level and their log2 fold changes in merged single-cell data that mimics the bulk level measurements. The results from edgeR are shown. **(b)** The expression of these genes at the single-cell level over the course of the infection in two cell types. Expression was calculated as log-transformed UMI counts normalized by the host library size. The violins below the dots are colored by samples. **(c)** Proportion of cells in which these genes are detected over the course of the infection in two cell types. The colors are consistent with those in panel (b).

To evaluate cell-to-cell variation in the host response manifested by the alteration in the host transcriptome following IAV infection, we performed un-supervised cell clustering solely using the host transcriptional data. We then annotated single cells in each subpopulation with information about the relative abundance of viral transcripts and the DVG/FL ratio. We observed a correlative pattern between the distribution of the relative abundance of virus transcripts and the host transcriptional landscape across cells within a population, as cells with a high abundance of virus transcripts formed a separate cluster by 12hpi in both cell types and this population structure, marked by a drastic difference in the level of viral transcripts between clusters, was retained at 24hpi (Fig. 3a and Fig. 3b). However, cell-cycle status, which contributes to the heterogeneity of the cell population by 12hpi and the formation of clusters in distinct cell-cycle stages (Supplementary Fig. 13a and Supplementary Fig. 13b) with a similar abundance of virus transcripts, does not seem to have a significant impact on population structure in A549 cells at 24hpi. Consistent with previous reports on G0/G1 cell-cycle arrest [24, 25], at 24hpi we observed a drastic increase in the proportion of cells in the G0/G1 phase as compared to earlier time-points (Supplementary Fig. 13c). Interestingly, this same pattern was not observed in HBEpC cells at the same time-point; in these cells the host and viral transcriptome landscape at 24hpi is similar to that in A549 cells at 12hpi, suggesting a temporal difference in host response and viral replication between cell types.

**Figure 3.**
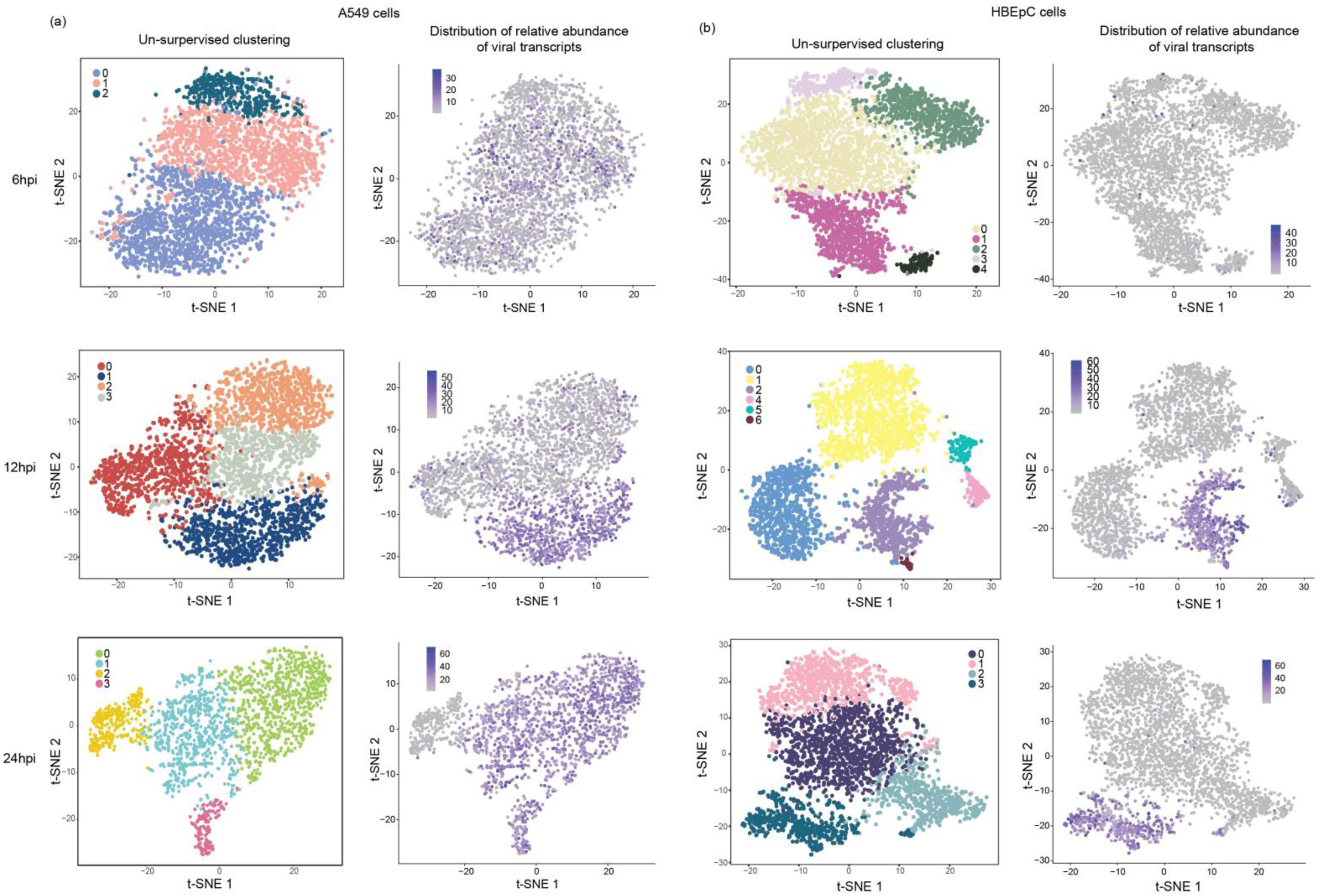
Comparison of cell clustering patterns and the distribution of viral transcript relative abundance. PR8-infected **(a)** A549 and **(b)** HBEpC cells harvested at each time point were subjected to un-supervised clustering based on the host transcriptome and visualized on a t-SNE plot. Each dot on a t-SNE plot represents a cell. Cell clustering patterns, in which cells were colored by their cluster identity, are shown in the left panels, and the distribution of the relative abundance of viral transcripts at different time points in individual cells is shown in the right panels.

To characterize host genes driving the changes associated with viral transcription, we assessed the over-representation of Gene Ontology (GO) terms associated with differentially expressed genes in each subpopulation of A549 and HBEpC cells compared to the rest of the cells at 24hpi. In two subpopulations of A549 cells with a high relative abundance of virus transcripts (clusters 0 and 1 in Fig. 4a), there is an enrichment of genes involved in viral transcription, RNA processing, translation, SRP-dependent co-translational protein targeting to the membrane, and mitochondrial electron transport (Fig. 4a). These GO terms are also associated with genes highly expressed in HBEpC cells with the highest abundance of virus transcripts at 24hpi (cluster 3 in Fig. 4b; p < 2.2 x 10^-16^ compared to the other clusters). In contrast, genes involved in antiviral responses, such as the type I IFN signaling pathway and the negative regulation of viral genome replication, are highly expressed in the other two subpopulations of A549 cells with a similar or severely reduced relative abundance of virus transcripts (clusters 3 and 2, respectively in Fig. 4a). However, some antiviral genes are differentially induced in cells with different levels of viral transcription. For example, type I and III IFNs, as well as a subset of ISGs, are highly expressed in cluster 3, and another subset of ISGs are highly expressed in cluster 2. In HBEpC cells, the antiviral responses, such as the type I IFN signaling pathway, are primarily observed in a cluster that has a low abundance of virus transcripts (cluster 2 in Fig. 4b) and is mostly comprised of cells in the G0/G1 phase (Supplementary Fig. 13b at 24hpi), as seen for A549 cells at 12hpi (Supplementary Fig. 13a and Supplementary Fig. 14).

**Figure 4.**
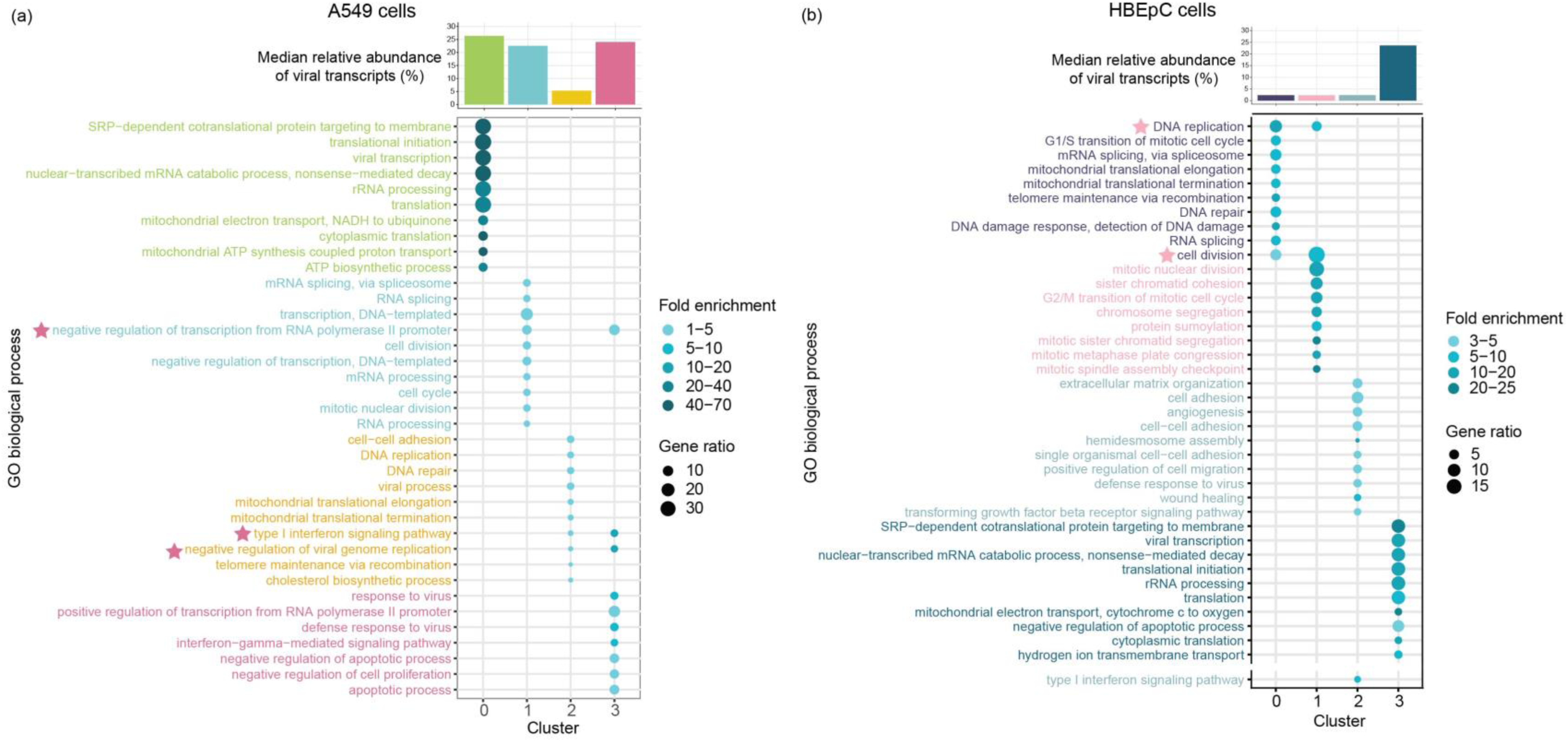
Enriched biological process gene ontology (GO) terms in A549 and HBEpC cells at 24hpi. **(a-b)** Biological process GO terms associated with over-expressed genes in each cluster of (**a**) A549 and (**b**) HBEpC cells. The top 10 over-represented GO terms in each cluster are shown in the upper panel and another over-represented GO term related to type I IFN signaling pathway in clusters 2 of HBEpC cells are shown in the lower panel. Each GO term is denoted by a dot. The color intensity of each dot indicates the fold enrichment of the corresponding GO term and the size corresponds to the ratio of queried genes in the gene set associated with a given GO term. The GO terms enriched in each cluster are color-coded by clusters. The asterisks in the upper panel denote the GO terms enriched in two clusters and their colors indicate the identity of the other cluster besides the one denoted by the color of the GO term. The bar chart above the bubble chart of the GO terms shows the median relative abundance of the viral transcripts across cells in each cluster. The colors of the bar are consistent with that of the GO terms.

To further understand how viral and host transcriptional activities are coordinated in clusters of cells that are subjected to different levels of viral stimuli, we performed gene co-expression network analyses using multiscale embedded gene co-expression network analysis (MEGENA) [26] and subsequent correlation analyses to directly characterize virus-host interactions. This was done for each cluster individually for A549 and HBEpC cells collected at 24hpi to identify modules of co-expressed genes representing coherent functional pathways. The relevance of a module to viral infection was evaluated by correlating the first principal component of a module with the relative abundance of virus transcripts. Consistent with the processes identified in the GO term enrichment approach using differentially expressed genes identified within a given cluster of cells (Fig. 4), those related to the host cell cycle and viral life cycle, including transcription, translation, RNA processing, metabolic process, and protein localization to membrane, were enriched in the modules that were correlated with viral transcription and identified in the clusters of A549 cells with a high abundance of virus transcripts (Supplementary Fig. 15). Conversely, the host innate immune response (e.g., response to type I IFN) was associated with viral transcription-correlated modules in cells with the lowest abundance of virus transcripts (cluster 2 of A549 cells at 24hpi in Fig. 5a). For example, module M100 negatively correlated with relative abundance of virus transcripts (ρ = −0.262, p = 2.9 x 10^-5^) in cluster 2 of A549 cells has genes—including *ISG20, RNF213, IFI35, STAT1*, *RBCK1*, *IFI27*, *OAS2*, *OAS3*, *XAF1*, *STAT2*, *IFIT5—*that are involved in the innate antiviral response and that were significantly highly expressed, as compared to their expression in other cells of the same population (Fig. 5b). In HBEpC cells, viral transcription-correlated modules, such as M227, enriched for type I IFN signaling genes (e.g., *ISG15*, *MX1, MX2, IFI35, IFI27, IFIT1, IFITM3, OAS1, OASL, ISG20*.) and positively correlated with relative abundance of virus transcripts (ρ = 0.103, p = 0.021; Fig. 5c and 5d), were detected in one cluster of cells with a low abundance of virus transcripts (cluster 2 of HBEpC cells at 24hpi in Fig. 5c). Modules correlated with viral transcription in the other two clusters of HBEpC cells with a low abundance of virus transcripts (clusters 0 and 1 in Supplementary Fig. 16) were enriched for genes associated with the host cell cycle, consistent with observations in A549 cells at 12hpi (Supplementary Fig. 14), while the viral transcription correlated modules in HBEpC cells with a high abundance of virus transcripts (cluster 3 in Supplementary Fig. 16) were enriched for genes involved in the ER-associated ubiquitin dependent protein catabolic process, small molecule metabolic process, vesicle mediated transport, etc. Overall, by identifying modules of genes that have a direct correlation with virus transcript abundance, the gene co-expression network analysis further demonstrates a complex association between virus transcript abundance and host responses, consistent with the pattern revealed by GO term analysis of highly expressed genes in individual clusters, and thus provides additional evidence of complex virus-host interactions.

**Figure 5.**
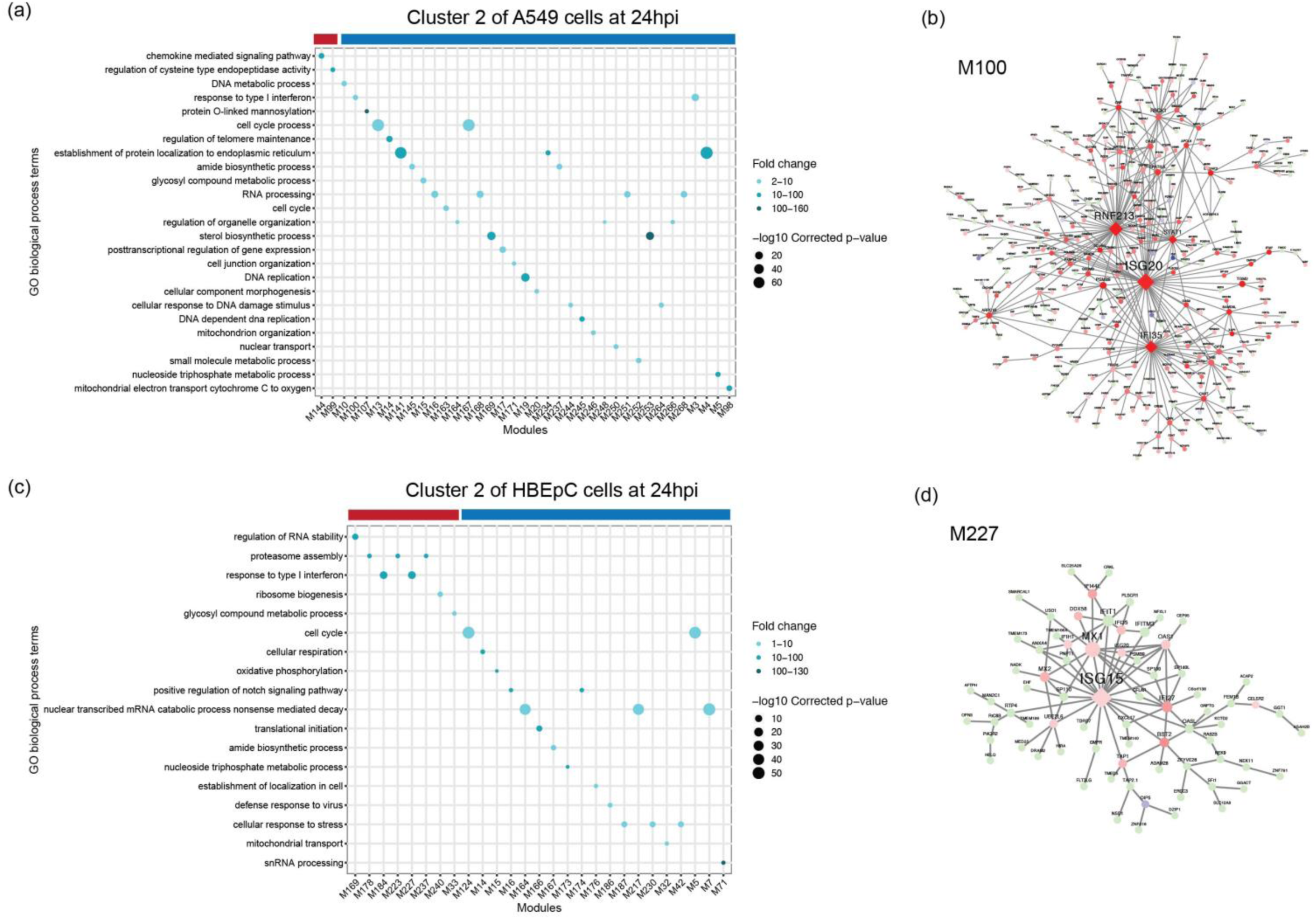
Functional MEGENA modules in clusters of A549 and HBEpC cells at 24hpi. Left panels show enriched biological process GO terms in the modules correlated with the relative abundance of virus transcripts in **(a)** cluster 2 of A549 cells and **(c)** cluster 2 of HBEpC cells at 24hpi. Each GO term is denoted by a dot. The color intensity of each dot indicates the fold enrichment of the corresponding GO term and the size corresponds to the -log10 transformed corrected p-value for a given GO term. Red and blue color-bars above the dot plot denote positive or negative correlation of viral transcription with corresponding modules, respectively. Right panels show the MEGENA network of modules enriched with the innate immune response, including **(b)** module M100 in cluster 2 of A549 cells and **(d)** module M227 in cluster 2 of HBEpC cells. Red and blue nodes represent significantly up- or down-regulated genes compared to the other cells in the population, while light-green nodes denote genes that are not significantly differentially expressed. Diamond nodes indicate key regulators. The size of the nodes indicates node strength after Multiscale Hub Analysis within the MEGENA pipeline. Modules that have a significant correlation with the relative abundance of virus transcripts and enriched GO biological process terms are shown in the GO term plots (**a** and **c**). Module member information for all viral transcription correlated-modules can be found on Github (https://github.com/GhedinLab/Single-Cell-IAV-infection-in-monolayer/tree/master/SupplementaryTables).

### Differential expression of host genes linked to defective viral genomes

Since DVGs are known to play an important role in IFN induction and inhibition of viral replication (reviewed in [20]), we determined whether the accumulation of DVGs was associated with attenuated viral transcription and induction of innate immune antiviral response genes. We observed a subtle but consistent inverse trend between the relative abundance of virus transcripts and the PB2 and PB1 DVG/FL ratios at 12hpi in A549 cells and at 24hpi in HBEpC cells (Fig. 6a-d). The cluster of cells with the highest abundance of virus transcripts (cluster 1 of A549 cells at 12hpi in Fig. 6a and cluster 3 of HBEpC cells at 24hpi in Fig. 6b) has the lowest level of PB2 and PB1 DVG/FL ratios (Fig. 6c-d; p < 2.2 x 10^-16^ comparing PB2 and PB1 DVG/FL ratios in cluster 1 of A549 cells or cluster 3 of HBEpC cells against the other clusters). Similarly, in A549 cells at 24hpi, we see the lowest abundance of virus transcripts in cells (cluster 2 in Fig. 6e) that have a higher level of PB2 DVG/FL ratios (cluster 2 in Fig. 6f; p < 2.2 x 10^-16^ compared to clusters 0 and 1) and the highest level of PB1 DVG/FL ratios (p = 1.943 x 10^-9^ comparing cluster 2 against the other clusters). Moreover, innate immune response genes are highly expressed in cells with either an elevated DVG/FL ratio for both DVG PB2 and PB1 transcripts and a low abundance of virus transcripts (cluster 2 of A549 cells at 24 hpi in Fig. 6e-f), or an elevated DVG/FL ratio for PB2 transcripts and a relatively high abundance of virus transcripts (cluster 3 in Fig. 6e-f; p = 4.036 x 10^-11^ compared to clusters 0 and 1 in Fig. 6f). It suggests that both DVG and FL transcripts could play a role in stimulating the innate immune response and that the enrichment of DVG transcripts in the total virus transcript pool is associated with strong immunostimulation, independent of viral transcription levels. Notably, the abundance of two dominant and ubiquitous PB2-DVG transcripts in the total virus transcript pool varies between cells and across clusters. The DVG PB2 transcripts carrying the same deletion sites as for PR8-DI222 were highly abundant in the cluster of cells with the lowest abundance of virus transcripts (cluster 2 of A549 cells at 24 hpi in Fig. 6g; p < 2.2 x 10^-16^ compared to the other clusters), while the other DVG PB2 transcripts corresponding to PR8-DI162 were highly abundant in both cluster 2 and cluster 3 (Fig. 6g; p = 2.067 x 10^-10^ and p = 0.001241, respectively, compared to clusters 0 and 1), suggesting a potential difference in the induction of innate immune response genes by different DVG species. We did not detect a difference in the relative abundance of defective PA transcripts among cells with different levels of viral transcription (Supplementary Fig. 17).

**Figure 6.**
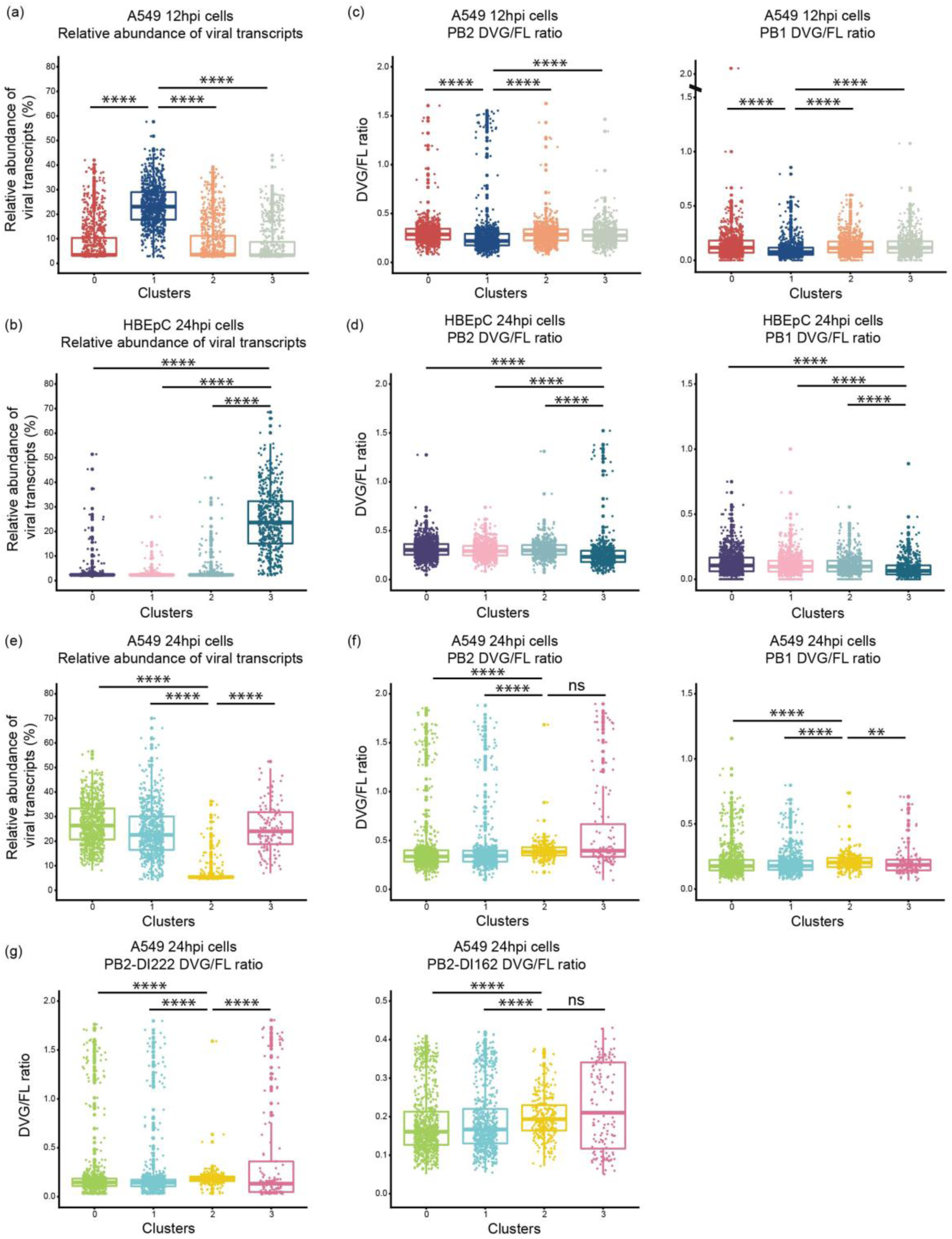
Distribution of the relative abundance of viral transcripts and DVG/FL ratio for DVG PB2 and PB1 transcripts in each cluster of cells. **(a, b, and e)** Box-plot of the relative abundance of viral transcripts in each cluster of **(a)** A549 cells at 12hpi, **(e)** 24hpi, and **(b)** HBEpC cells at 24hpi. **(c, d, and f)** The DVG/FL ratio for DVG PB2 and PB1 transcripts calculated as the ratio of gap-spanning reads to non-gap-spanning reads aligned to a 3’ coverage peak region derived from the corresponding viral segments in each cluster. **(g)** The DVG/FL ratio for DVG PB2 transcripts carrying deletion sites similar to those in PR8-DI222 and PR8-DI162 viruses in each cluster of A549 cells at 24hpi. All box plots show the first and third quantiles as the lower and upper hinges, the median in the center, and 1.5 * inter-quartile range (IQR) from the first and third quantiles as the whiskers. The significance levels of pairwise comparisons determined by one-tailed Wilcoxon rank sum test were denoted by the asterisks: * p ≤ 0.05, ** p ≤ 0.01, *** p ≤ 0.001, **** p ≤ 0.0001, and “ns” as not significant.

Given the enrichment of PB2 DVGs and the over-expression of ISGs in cluster 2 of A549 cells at 24hpi compared to the rest of the cells in the same population, we experimentally validated the immunostimulatory impact of PB2 DVGs on the host cells. We infected A549 cells with each of our 3 PR8-DIs (PR8-DI162, -DI222, or -DI244) or with WT virus, and quantified the expression level of the influenza virus M gene and of ISGs that were shown to be significantly highly expressed in cluster 2 of A549 cells, including *IFI35, IFI27*, and *MX1*, the type I (IFN beta) and III (IFN lambda) IFNs that were shown to be significantly highly expressed in cluster 3 of A549 cells (Supplementary Fig. 18a and 18b). Due to their limited ability to replicate without a helper virus, DI viruses cultured on their own exhibit a significantly lower level of viral transcription compared to WT virus (Supplementary Fig. 18c). Despite the lower replication of DIs, there was a higher induction of ISGs in DI-infected cells compared to cells infected with WT virus, while there was a comparable or marginally higher induction of IFN beta (IFN-ß) and IFN lambda (IFN-λ) in the WT-infected cells (Supplementary Fig. 18c). Together, these data demonstrate that at the late stage of infection PB2 DVGs have a strong immunostimulatory effect on the host cells manifested by a significant induction of ISGs even at very low levels of viral transcription and independent of type I and III IFN expression levels.

## DISCUSSION

The goal of this study was to quantitatively and qualitatively characterize viral and host factors that contribute to cell-to-cell transcriptional variation observed during IAV infection. We identified DVGs as potentially important factors in the temporal variation of the host cell transcriptome. The substantial cell-to-cell variation in viral replication and host response that we detected highlight the potential of single-cell virology to provide novel insight into complex virus-host interactions.

The inherent stochasticity of molecular processes, especially during the initial stages of infection, can contribute to the cell-to-cell variation in viral replication [1]. Evidence from studies examining low MOI infections with different virus strains and cell lines suggests that this intrinsic stochasticity can impact viral replication and result in the loss of viral genome segments, transcripts, or proteins [1, 8–10]. Similarly, at different time points of the high MOI infection, we observed cell-to-cell variation in viral transcription, including that in segment-specific transcription. Although the mechanisms leading to this observation are unclear based on our data, it may be explained by the following possibilities as discussed in [10], including (i) the absence of one or more viral genome segments in infecting virions, (ii) a deficiency in intracellular trafficking of incoming viral ribonucleoprotein complexes (vRNPs), (iii) the random degradation of vRNAs prior to transcription, or (iv) mutations resulting in decreased stability of viral mRNAs.

Another source of variability in viral replication stems from the emergence and accumulation of various vRNA species synthesized by the error-prone viral polymerase. Indeed, viral genetic diversity, including amino acid mutations in the NS1 and PB1 proteins, the absence of the NS gene, and an internal deletion in the PB1 gene, contributes to the cell-to-cell variation in the innate immune response as shown in a recent report on a low MOI infection [27]. Although the 3’ sequencing strategy we used in our study limits the identification of mutations in the full-length segments, our experimental setup provides a unique opportunity to evaluate the diversity of DVGs by characterizing the DVG transcripts, since a high MOI infection promotes the accumulation of DVGs. Late stages of infection also emphasize the impact of accumulated DVGs on the host response. We identified a diverse pool of DVG transcripts derived from the three polymerase segments, including several dominant species. Notably, the fact that all the dominant DVG PB2 and PB1 transcripts identified in the infected cells were also present in the virus stock, and that the dominant DVG PB1 transcripts were detected in an increasing proportion of cells over the course of infection, suggests potential cellular transmission of DI viruses carrying these DVGs. However, we cannot exclude the possibility that the regions where the hot spots were located are most prone to polymerase skipping and that DVG enrichment over the course of the infection resulted in their abundance above a detection threshold in an increasing number of cells.

The accumulation of DVGs can have consequences on viral replication by directly competing with the full-length genomes and modulating host innate immune responses. The immune-modulatory effect of DVGs has been attributed to efficient activation of the IFN induction cascade and antiviral immunity, a mechanism well-known for the non-segmented negative-sense RNA viruses, such as the murine parainfluenza virus Sendai (SeV) (reviewed in [20]). Consistent with previous reports, we detected enrichment of highly expressed genes involved in the IFN response in one subpopulation of cells with significantly attenuated viral transcription. In this cluster of cells we also observed that DVG transcripts accumulated to significantly higher levels compared to FL transcripts. The co-expression network analysis provides an opportunity to identify modules of co-expressed host genes and directly correlate them with the level of viral transcription revealing groups of innate response genes that are potentially differentially expressed under the effect of viral transcription. The observation was further validated in our DI- or WT-infection assay, in which we detected a significantly higher expression of some ISGs in DI-infected cells independent of the level of viral transcription and that of IFNs at the late stage of the infection. The IFN mRNA dose-independent ISG expression in DI-infected cells suggests a potential novel mechanism of DVG immunostimulation that is independent of IFN induction. Expression of a subset of ISGs can be induced in the absence of IFN signaling, as shown in other studies [28, 29]. It is likely to be at least partially attributed to similar and overlapping consensus DNA-binding motifs for the interferon regulatory factor (IRF) family, most notably IRF3 and IRF7 that are key regulators of type I IFN production, and for IFN-stimulated gene factor 3 (ISGF3) that promotes the expression of many ISGs in response to IFN signaling [30–33]. For example, a IRF3 ChIP-seq experiment with adenovirus-infected human primary bronchial/tracheal epithelial cells demonstrated IRF3 binding to the promoter region of induced ISGs, including MX1 and ISG15, which have the IFN-stimulated response element (ISRE) bound by IRFs and ISGF3 in the promoter regions [34]. Further investigation of ISG induction in the absence of IFN signaling during DI infection will elucidate this potential mechanism of DI immunostimulation. In addition, given the fact that some ISGs could be highly expressed in cells with a high level of viral transcription or a high abundance of DVGs suggests that other viral factors, besides the accumulation of DVGs, may also play an important role in triggering the innate immune response. Nevertheless, unlike SeV DVGs that form “copy-back” structures due to complementary termini generated by “copying back” the authentic 5’ terminus at the 3’ terminus [35, 36], IAV DVGs share identical termini with the full-length genomes. In the copy-back SeV DVGs, a stretch of dsRNA adjacent to a 5’-triphosphate serves as an effective RIG-I ligand [20]; however, the mechanism of IAV DVGs underlying a more effective immunostimulation compared to the full-length genomes remains elusive [20]. A proposed hypothesis is that the unencapsidated replication products of small DVGs can potentially activate RIG-I if they can reach the cytoplasm [20], as observed *in vitro* and *in vivo* where short influenza RNA templates can be replicated in the absence of nucleoprotein (NP) [37]. In addition, the interfering and immune-modulatory abilities of DVGs could also be attributed to DVG-encoded proteins that directly interact with mitochondrial antiviral-signaling (MAVS) and act independently of RIG-I in IFN induction, as previously reported [38].

Our approach enabled us to characterize and quantify the abundance of diverse DVG species that accumulated in individual cells. It also allowed us to associate the enrichment of certain DVG species with the attenuation of viral transcription and the induction of innate immune responses at single-cell resolution. For example, the accumulation of PB2-DVGs, especially including one type of dominant PB2-DVG species (PR8-DI222), in one subpopulation of cells was associated with significantly attenuated viral transcription and a strong host innate immune response manifested by high expression of some ISGs, although we cannot rule out the effect of other non-dominant defective PB2-DVGs, DVGs derived from other polymerase segments, or certain mutations in stimulating host responses. Diverse DVG species can play different roles in shaping the virus-host interaction. However, further investigation is necessary to elucidate the dynamics and impact of DVG species distribution during pathogenesis, enhancing the understanding of the relationship between DVG dynamics and infection outcome. Our characterization of DVGs and corresponding host responses reveals complex virus-host interactions and an underappreciated single-cell level of immune-modulation by DVGs. The complex gene expression pattern underlying the host innate immune response demonstrates the intricate effects of various viral factors, including—but probably not limited to—the accumulation of DVGs, on shaping infection outcome at the single-cell level.

## MATERIALS AND METHODS

### Cells and virus

Human lung adenocarcinoma epithelial A549 cells (ATCC, Virginia, USA) were maintained in Kaighn’s modified Ham’s F-12 medium (F-12K) supplemented with 10% fetal bovine serum (FBS). Primary human bronchial epithelial cells (HBEpC) (PromoCell, Heidelberg, Germany) were maintained in PromoCell airway epithelial cell growth media with SupplementMix (PromoCell). Madin-Darby canine kidney (MDCK) cells were maintained in minimum essential medium (MEM) supplemented with 5% FBS. Human embryonic kidney epithelial 293T cells (ATCC, Virginia, USA) were maintained in Dulbecco’s Modified Eagle Medium (DMEM) supplemented with 10% FBS. PB2 protein-expressing modified MDCK (AX4/PB2) [39] cells were maintained in MEM supplemented with 5% Newborn Calf Serum (NBCS), 2 µg/ml puromycin and 1 µg/ml blasticidin. 293T and AX4/PB2 cells were co-cultured with DMEM supplemented with 10% FBS. Viral strain A/Puerto Rico/8/34 (H1N1) was plaque purified and propagated in MDCK cells. Viral titers were determined by plaque assay in MDCK cells and sequences confirmed by Illumina MiSeq sequencing.

### Infection assays

Subconfluent monolayers of A549 cells in T-25 flasks were washed with A549 infection media (F-12K supplemented with 0.15% bovine serum albumin (BSA) fraction V and 1% antibiotic-antimycotic) and infected with influenza virus at a multiplicity of infection (MOI) of 5 in 0.5 mL pre-cooled A549 infection media. The flasks were incubated at 4°C for 30 minutes with agitation every 5-10 minutes followed by addition of 4.5 mL pre-warmed A549 infection media and transferred to a 37°C, 5% CO_2_ incubator for further incubation. HBEpC cells were infected similarly except the HBEpC infection media was PromoCell airway epithelial cell growth media. The inoculum was back-titrated to confirm the desired MOI was used. The virus-infected cells were collected at 6, 12, and 24 hours post infection (hpi) and the mock-infected cells were collected at 12 hpi. There was no substantial cell death observed at any time point. Cells were extensively washed before re-suspension in PBS containing 0.04% BSA. A proportion of cells from one of the two duplicate flasks per time point were subject to 10X Genomics Chromium single-cell library preparation. The remaining cells in the two flasks were used for conventional bulk RNA-seq library preparation. Notably, due to a failure in the single-cell library preparation for the PR8-infected HBEpC cells collected at 12hpi, we replaced this sample by repeating the same infection assay with HBEpC cells.

### Sample preparation, library construction, and sequencing

10X Genomics single cell library preparation with Chromium 3’ v2 chemistry was performed following the manufacturer’s protocol. A range of 3,900-7,500 cells were used as input into each single cell preparation, with a median of ∼3,353 cells (range 2,075-5,254) obtained following sequencing, as described below. Sequencing was performed on the HiSeq 2500 in HighOutput mode (v2) with one library per lane following the manufacturer’s recommended sequencing configuration (i.e., paired-end read 1: 27 bp, read 2: 99 bp, and i7 index: 8 bp). An average of 59,200 reads per cell were obtained. For bulk mRNA sequencing, total RNA from two replicate flasks per time point was extracted using RNeasy Mini Kit (Qiagen, Hilden, Germany) with on-column DNA digestion using RNase-free DNase (Qiagen). The conventional bulk RNA-seq library preparation was performed using NEBNext Ultra II RNA library prep kit for Illumina following poly(A) mRNA enrichment. Libraries were multiplexed and sequenced on an Illumina NextSeq 500 in HighOutput 2×75 bp mode (v3). Viral genomic RNA (vRNA) of the virus stocks used in the infection assay was amplified using multi-segment reverse-transcription PCR (M-RTPCR) [40]. The M-RTPCR product was subject to Illumina library preparation with Nextera DNA library prep kit and sequenced on the HiSeq 2500 in Rapid 2×150 bp mode (v2).

### Upstream computational analyses of single-cell RNA-seq data

The Cell Ranger (v2.1.0) single cell software suite (10X Genomics) was applied to the single-cell RNA-seq data to perform the alignment to the concatenated human (hg19) and influenza A virus (A/Puerto Rico/8/34) reference with STAR (v2.5.3a) [41] and gene counting with the processed UMIs for each cell. The UMI counts for the viral reads with the CIGAR string containing a gap (i.e., “MNM”) and different UMI sequences comparing to the ungapped reads were added into the expression matrix later, since they were not considered during gene counting. Given the fact that we performed 3’-end sequencing, transcripts derived from alternative splicing of M and NS segments, encoding M1/M2 and NS1/NEP respectively, could not be distinguished in our analyses, as those transcribed from the same genes share the identical 3’-end.

To identify and exclude low quality cells in the single cell data from the downstream analyses, the following metrics were calculated for each sample and were employed to filter out cells that failed to meet these criteria: 1) remove cells with fewer than 2,000 detected genes; 2) remove cells with an alignment rate less than the mean minus 3 standard deviations; 3) remove cells with a number of reads after log10 transformation not within 3 standard deviations below or above the mean; 4) remove cells with the number of UMI after log10 transformation not within 3 standard deviations below or above the mean; 5) remove cells with the percentage of reads aligned to mitochondrial genes not within 3 standard deviations below or above the mean; 6) remove mock-infected cells with 1 viral read detected. Although we filtered cells based on the size of the total mRNA pool (i.e., total reads) as one of the criteria described above, we also checked the distribution of the size of the host mRNA pool (i.e., host reads) for individual cells. As reported in recent virus-infected single cell studies [9, 42], while virus-induced host shutoff of gene expression [43] is likely to be mainly mediated by host mRNA degradation, it remains unclear what effect viral transcription has on the size of the total mRNA pools. In our study, we observed that virtually all the cells with substantially smaller or larger host mRNA pools–where the number of host reads after log10 transformation was not within 3 standard deviations below or above the mean–were already being excluded when applying the 6 criteria described above, thus we decided to filter cells based on these. Approximately 120-250 cells were eliminated from each sample, with a median of 164 cells per sample. We further removed genes that were detected in fewer than three cells. After initial cell and gene quality control, the majority of samples had approximately 3,000-3,500 cells left for downstream analysis. The expression levels of host genes were normalized based on the size of the host mRNA pool.

### Cell clustering with Seurat

An unsupervised cell clustering to identify subtle changes in the population structure after infection was performed on each time point data separately following the procedures of the Seurat package (v2.1.0) [44] using the normalized, scaled, and centered host gene expression matrix, without considering the relative abundance of virus transcripts within individual cells. Briefly, the highly variable genes with average expression < 4 and dispersion > 1 were used as input for the PCA. The statistically significant and biological meaningful PCs determined by the built-in jackstraw and elbow analyses and manually exploration were retained for visualization by t-distributed stochastic neighbor (t-SNE) and subsequent clustering by a shared nearest neighbor (SNN) graph-based approach. The legitimacy of the initially identified clusters was validated using the “ValidateClusters” function in Seurat, which built a support vector machine (SVM) classifier with significant PCs and then applied the accuracy cutoffs of 0.9 and the minimal connectivity threshold of 0.001. To identify markers that are differentially expressed among clusters, the “FindMarkers” function in Seurat was used with the different test options, including “bimod” [45], “poisson”, “negbinom”, and “MAST” (v1.4.1) [46]. Only the genes that showed at least 0.25-fold difference on the log-scale between two groups and were expressed in at least 25% of the cells in either group were tested for differential expression. Significantly differentially expressed genes for each cluster (i.e., the markers) were identified by applying the adjusted p-value cut-off of 0.05. Markers identified by all four methods were retained. GO term over-representation analysis of the up-regulated markers was performed with the online service DAVID [47, 48] by applying the Benjamini p-value cut-off of 0.05. To determine the cell cycle stage associated with individual cells, cell-cycle scoring and assignment were performed with Seurat using the “CellCycleScoring” function based on the expression of canonical markers [49].

### Computational analyses of bulk RNA-seq and virus stock sequencing data

The raw bulk RNA-seq and virus stock sequencing data was first trimmed with trimmomatic (v0.36) [50] to remove the adaptors and trim off low quality bases. Reads with a minimal length of 36 bases in the trimmed bulk RNA-seq dataset were aligned to the concatenated human (hg19) and influenza A/Puerto Rico/8/34 (H1N1) reference with STAR (v2.5.3a) [41] with the default parameters, and counted with featureCounts [51] in the Subread package (v1.5.1) [52], while reads in the virus stock sequencing dataset were aligned to the influenza A/Puerto Rico/8/34 (H1N1) reference with STAR (v2.5.3a) [41]. Differential expression analysis was performed with DESeq2 (v1.18.1) [53] and edgeR (v3.20.5) [54, 55] using the bulk RNA-seq and the merged single-cell RNA-seq data. Host genes with the adjusted p-value < 0.05 were identified as significantly differentially expressed at each time point.

### Deletion junction identification, filtering, and quantification

Reads that aligned to both ends of a viral segment, with each aligned portion comprised of at least 10 nucleotides in length, were collected. Following UMI de-duplication, reads with junction coordinates within a 10-nucleotide window were grouped together. We excluded from downstream analyses reads with junction coordinates that occurred fewer than 10 times in each sample. To compare quantitatively gap-spanning reads for each segment across cells and samples, the number of gap-spanning reads was normalized to the total number of non-gap-spanning reads aligned to a 100nt region centering the coverage peak at the 3’ end. For the polymerase segments, gap-spanning reads were also filtered by the size of deletions (i.e. ≥ 1000 nucleotides). The 1000 nucleotides threshold for the polymerase segments was chosen based on the distribution of the size of the deletions (Supplementary Fig. 19), as the majority of the deletions are larger than 1000 nucleotides. As cells with low infection (especially those harvested at the early stage of infection or inoculated at a low MOI) typically have poor coverage for the viral segments of interest, the DVG/FL ratio in those cells calculated as described above is typically inflated and may even fail to be calculated because all the viral reads corresponding to a given segment are gap-spanning or there are no viral reads. To mitigate this effect, we overwrote those values to 0 in the datasets, including the dataset collected from HBEpC cells at 6hpi.

### Generation of PR8-derived DI162, DI222, DI244, and PB2del virus

To generate the PR8 PB2-DI162, -DI222, and -DI244 reverse genetics plasmids, gBlocks Gene Fragments (Integrated DNA Technologies, California, USA) were ordered using the corresponding defective PB2 genomic sequences identified in this study (PR8-DI162 and PR8-DI222) or previously reported (DI244) [18]. To generate the PR8 PB2del plasmid, a point mutation at nucleotide position 265 was introduced into the PR8 PB2 genomic sequence using the mutagenesis primer PR8-PB2-stopdel-F (5’-TTCATTTACTCCATTAAGTTTGTCCTTGCTC-3’). The reverse genetics plasmids were cloned into the pBZ61A18 reverse genetics vector as previously described [56]. To rescue the PR8 DI viruses, each of the sequence-confirmed PB2-DI or PB2del plasmids was co-transfected into 293T-AX4/PB2 co-cultured cells with the 7 reverse genetics plasmids for PR8 PB1, PA, HA, NP, NA, M, and NS and a PB2 protein-expression plasmid using Lipofectamine™ 3000 Transfection Reagent (Invitrogen, California, USA). The supernatant was collected on day 2 post-infection and passaged twice in the AX4/PB2 cell line, which expresses the PB2 protein in *trans* [39]. Viral titers were determined by TCID50 assay in AX4/PB2 cells.

### Interference test for PR8-DI162 and PR8-DI222 viruses and validation of their immunostimulatory effects

To test the interfering ability of PR8-DI162 and PR8-DI222 viruses that carry the deletions identified in two dominant PB2-DVG species from the single cell dataset, their inhibitory effects on wild type PR8 virus replication was quantified in cultured cells at different DI/WT ratios. Subconfluent monolayers of A549 cells in 12-well plates were washed with infection media (MEM supplemented with 0.15% BSA fraction V, 1% antibiotic-antimycotic and 1 µg/mL TPCK-treated trypsin) and each well was co-infected with one type of defective virus at a MOI of 5 and the PR8 virus either at a MOI of 0.005 for a DI/WT ratio of 1000:1 or at a MOI of 5 for a DI/WT ratio of 1:1 in 1 mL infection media. For a DI/WT ratio of 1000:1 co-infection, plates were transferred to a 37°C, 5% CO_2_ incubator and supernatants were collected at 2 hours post infection, and 1, 2, and 3 days post infection. For a DI/WT ratio of 1:1 co-infection, infection was similar to that used for single-cell RNA-seq, including incubating at 4°C for 30 minutes with agitation every 5-10 minutes after infection followed by adding pre-warmed A549 infection media and transferred to a 37°C, 5% CO_2_ incubator for further incubation. Supernatants were collected at 2, 6, 12, 24 hours post infection and titrated as described above. Wild type PR8 yield from collected supernatants was determined by TCID50 assay using MDCK cells, which do not support replication of DI viruses due to the lack of functional PB2 proteins.

To determine if the DI viruses could stimulate a higher expression of some ISGs identified in the network analysis, subconfluent monolayers of A549 cells in 12-well plates were infected with each DI virus (PR8-DI162, -DI222, or -DI244) or WT virus at a MOI of 10 in triplicate. Total RNA from cells collected at 24hpi was extracted using RNeasy Mini Kit (Qiagen, Hilden, Germany). 100ng RNA was subsequently used as template for reverse transcription using SuperScript IV system (Invitrogen, California, USA) with Oligo d(T)_20_ primer. qPCR was performed using PowerUp SYBR Green (Applied Biosystems, Massachusetts, USA) in triplicate with the following primers: IFI35: Hs.PT.58.38490206 (IDT, Iowa, USA); IFI27: IFI27fw (5’-GCCTCTGGCTCTGCCGTAGTT-3’) and IFI27rev (5’-ATGGAGGACGAGGCGATTCC-3’) [57]; MX1: MX1fw (5’-CTTTCCAGTCCAGCTCGGCA-3’) and MX1rev (5’-AGCTGCTGGCCGTACGTCTG-3’) [57]; IFNB: IFNBfw (5’-TCTGGCACAACAGGTAGTAGGC-3’) and IFNBrev (5’-GAGAAGCACAACAGGAGAGCAA-3’) [58]; IFNL: IFNLfw (5’-GCCCCCAAAAAGGAGTCCG-3’) and IFNLrev (5’-AGGTTCCCATCGGCCACATA-3’) [58]; β-actin (ACTB): ACTBfw (5’-GAACGGTGAAGGTGACAG-3’) and ACTBrev (5’-TTTAGGATGGCAAGGGACT-3’) [59]; FLU: mCDC-FluA-F-PR8 (5’-GACCAATCCTGTCACCTCTGA-3’) and mCDC-FluA-R (5’-AGGGCATTTTGGACAAAGCGTCTA-3’). For all the targets, qPCR parameters according to manufacturer’s recommendation were: 95°C for 10 min and then 45 cycles of 95°C for 15 s, 57°C for 15 s, and 60°C for 60 s. Fold change of target gene expression was calculated using the 2^−ΔΔ*C_T_*^ method normalized to ACTB.

### Gene co-expression network analysis

Multiscale Embedded Gene Co-Expression Network Analysis (MEGENA) [26] was performed to identify host modules of highly co-expressed genes in influenza infection. The MEGENA workflow comprises 4 major steps: 1) Fast Planar Filtered Network construction (FPFNC), 2) Multiscale Clustering Analysis (MCA), 3) Multiscale Hub Analysis (MHA), 4) and Cluster-Trait Association Analysis (CTA). A cutoff of 0.05 after perturbation-based FDR calculation was used. The differentially expressed genes are identified between virus-infected cells of a particular cluster and the rest of the cells in the same population determined by a t-test. Only the genes that showed at least 0.25-fold difference on the log-scale between two groups, and were expressed in at least 25% of the cells in either group, were tested for differential expression. Significantly differentially expressed genes were identified by applying the adjusted p-value cut-off of 0.05.

### Identification of enriched GO terms, key regulators in the host module, relative abundance of virus transcripts associated with host modules

To functionally annotate gene signatures and gene modules identified in this study, enrichment analysis was performed of the established gene ontology (GO) categories. The hub genes in each subnetwork were identified using the adopted Fisher’s inverse Chi-square approach in MEGENA; Bonferroni-corrected p-values smaller than 0.05 were set as the threshold to identify significant hubs.

The relative abundance of virus transcripts and the DVGs associated with the host modules were identified using Spearman correlation between the first principal component of the gene expression in the corresponding module and the relative abundance of viral transcripts. Significantly associated traits were identified using the Benjamini-Hochberg FDR-corrected p-value 0.05 as the cutoff.

## STATISTICAL ANALYSIS

The statistical significance of the changes in the relative abundance of viral and DVG transcripts between 3 or more groups of cells was first determined by the one- or two-tailed Kruskal-Wallis rank sum test, followed by the one-tailed Wilcoxon rank sum test to calculate the pairwise comparisons. The statistical significance of expression fold changes in the qPCR validation assay was determined using the two-tailed Student’s *t*-test. A p-value of ≤ 0.05 was considered statistically significant.

## CODE AVAILABILITY

The code used to generate all the results is available on Github (https://github.com/GhedinLab/Single-Cell-IAV-infection-in-monolayer).

## DATA AVAILABILITY

Sequencing data that support the findings of this study have been deposited in the Gene Expression Omnibus (GEO) repository with the accession codes GSE118773 (currently private record).

## ACKNOWLEDGEMENTS

We thank members of the Ghedin laboratory for feedback and discussion. We thank Tara Rock, Nicholas Rouillard, Olivia Micci-Smith, and Mohammed Khalfan of the Genomics Core Facility at the Center for Genomics and Systems Biology, New York University for sequencing. We thank Dr. Yoshihiro Kawaoka for providing the AX4/PB2 cells and Dr. Peter Palese and Dr. Adolfo Garcia-Sastre for providing the PR8 RGs plasmids. We thank Dr. Meike Dittmann for her feedback on IFNs and ISGs. This work was supported by NIAID/NIH U01 AI111598. This work was also supported in part through the NYU IT High Performance Computing resources, services, and staff expertise.

## AUTHOR CONTRIBUTIONS

All authors read and approved the manuscript. E.G. conceived and designed the experiments, supervised research, and wrote the manuscript. B. Zhou conceived and designed the experiments, supervised research, performed the infection assays, and wrote the manuscript. C.W. performed the infection assays and bulk RNA-seq library preparation, analyzed the data, and wrote the manuscript. C.V.F. performed the gene co-expression network reconstruction and module-based functional enrichment analysis and wrote the manuscript. T.C. performed the defective virus generation, the interference and validation assays, and wrote the manuscript. A.G. performed the sequence library preparation and contributed to the analyses. M.W. and B. Zhang contributed to data analyses. W.H., M.S., and R.S. performed the single-cell library preparation and contributed to the analyses.

## SUPPORTING INFORMATION

**Supplementary Figure 1.**
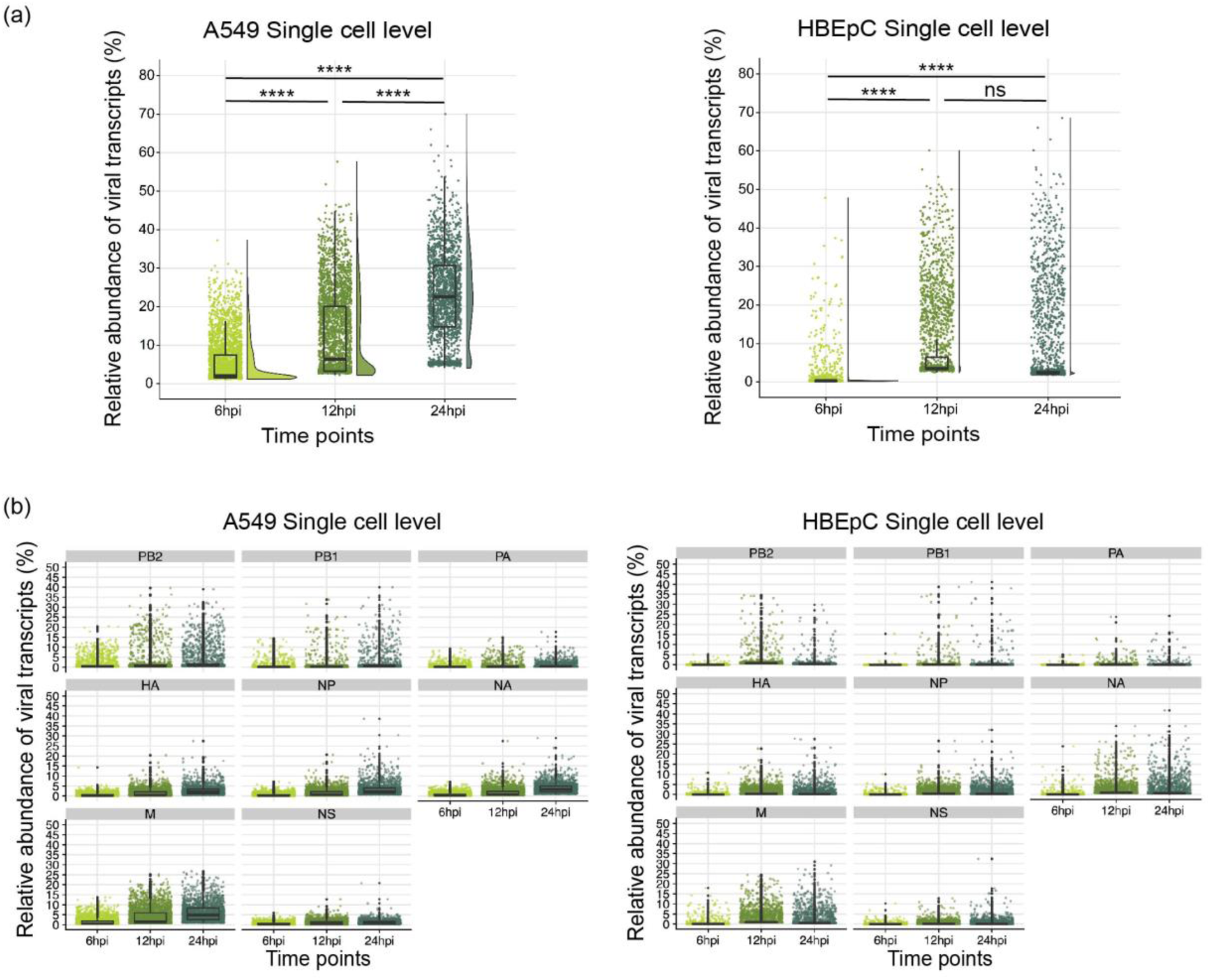
Distribution of the relative abundance of viral transcripts within individual cells over the course of the infection. **(a)** The overall relative abundance of viral transcripts was calculated as the percentage of viral reads in the pool of all the reads for each cell. The dots are colored by the time points. All pairwise comparisons were performed with one-tailed Wilcoxon rank sum test, for which the null hypothesis was that cells harvested at an earlier time point have a lower median relative abundance of viral transcripts than those harvested at a later time point. The significance levels were denoted by the asterisks: * p ≤ 0.05, ** p ≤ 0.01, *** p ≤ 0.001, **** p ≤ 0.0001, and “ns” as not significant. The data for HBEpC cells at 12hpi was collected from a repeated infection assay, due to the initial failure of single-cell library preparation for the corresponding sample. **(b)** Distribution of the relative abundance of viral transcripts derived from each segment within individual cells over the course of the infection. All box plots show the first and third quantiles as the lower and upper hinges, the median in the center, and 1.5 * inter-quartile range (IQR) from the first and third quantiles as the whiskers.

**Supplementary Figure 2.**
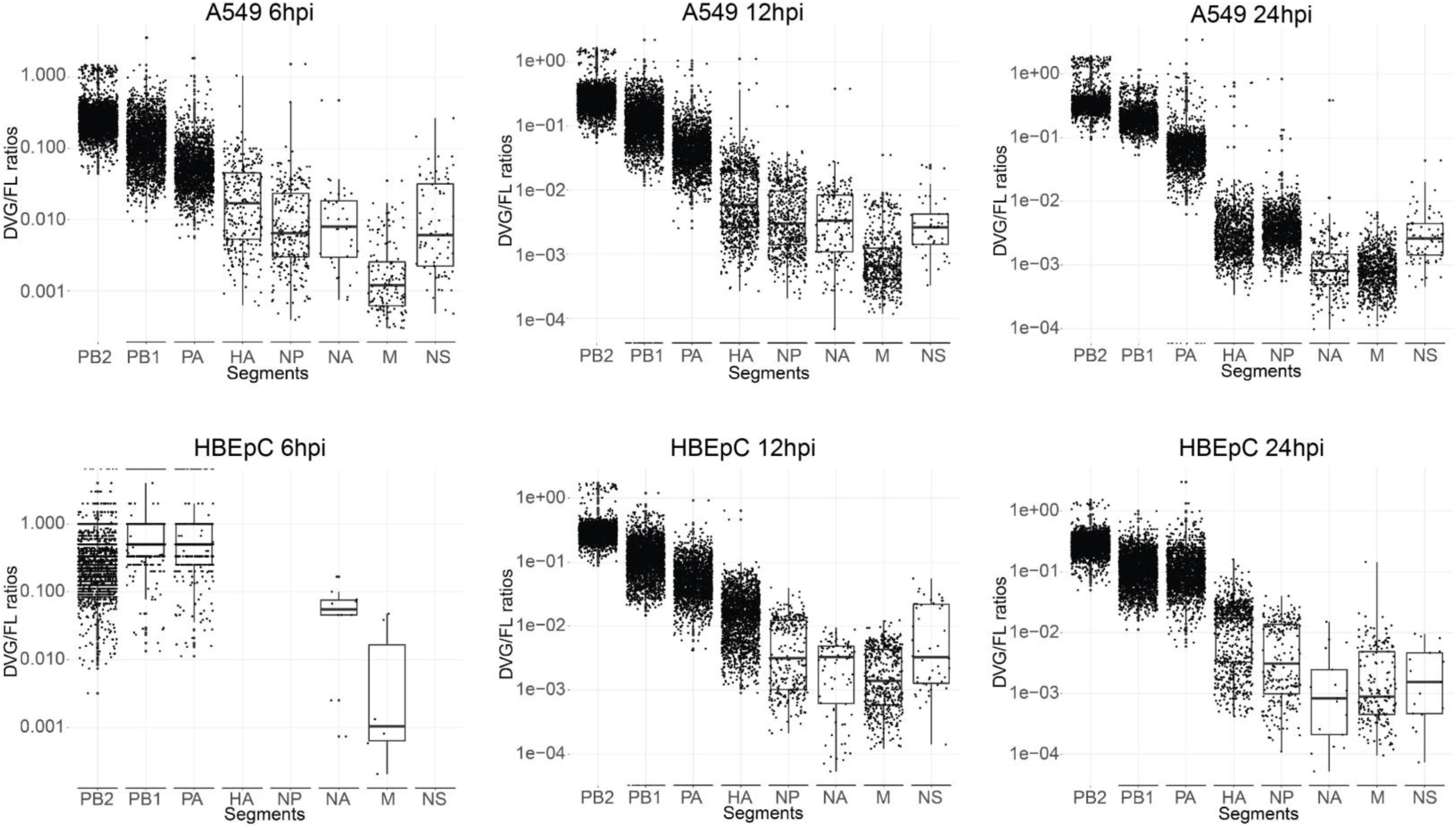
Distribution of DVG/FL ratios for each viral segment in two cell types over the course of the infection. DVG transcripts represented by gap-spanning reads were grouped by their junction coordinates and filtered by their frequency in each sample (i.e., >= 10 times)

**Supplementary Figure 3.**
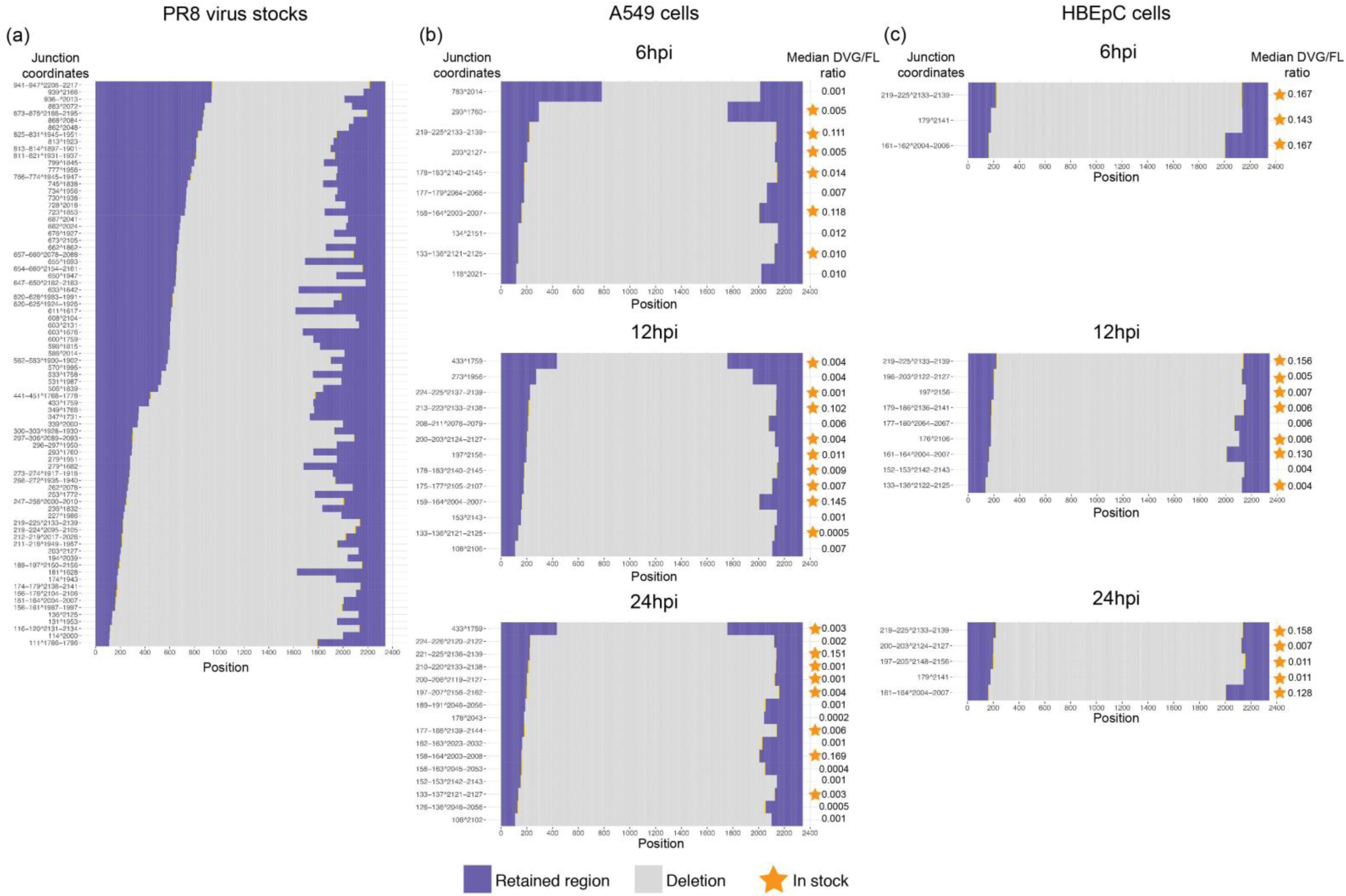
Distribution of junction sites in the PB2 segment and the medians of their DVG/FL ratio in cells where a given defective viral transcript was detected. **(a)** The defective PB2 segments detected in the PR8 virus stock used for infection. **(b-c)** The DVG PB2 transcripts detected in **(b)** A549 and **(c)** HBEpC cells over the course of the infection. Transcripts carrying the same junction sites as seen in the virus stock were denoted by asterisks.

**Supplementary Figure 4.**
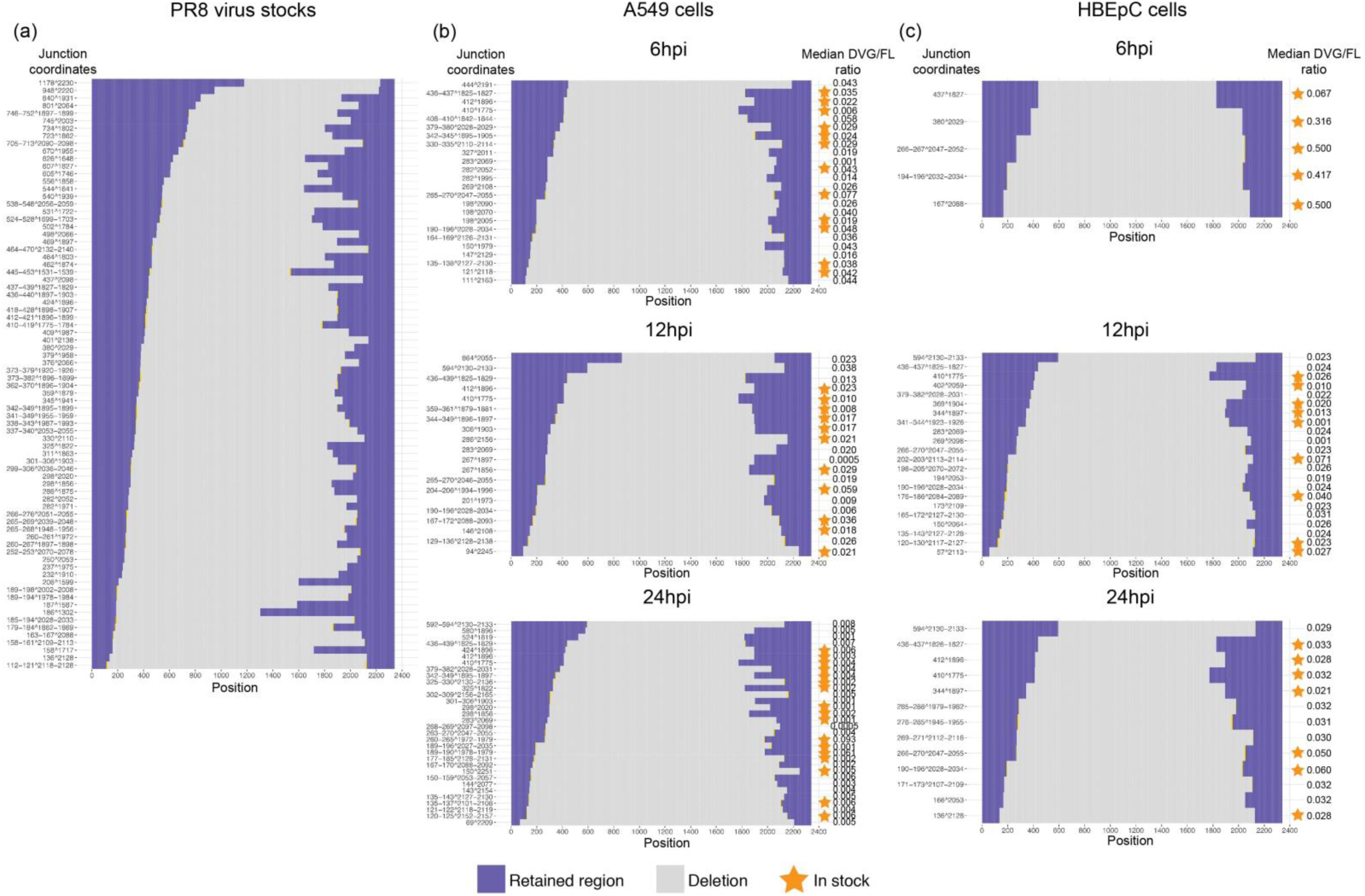
Distribution of junction sites in the PB1 segment and the medians of their DVG/FL ratio in cells where a given defective viral transcript was detected.

**Supplementary Figure 5.**
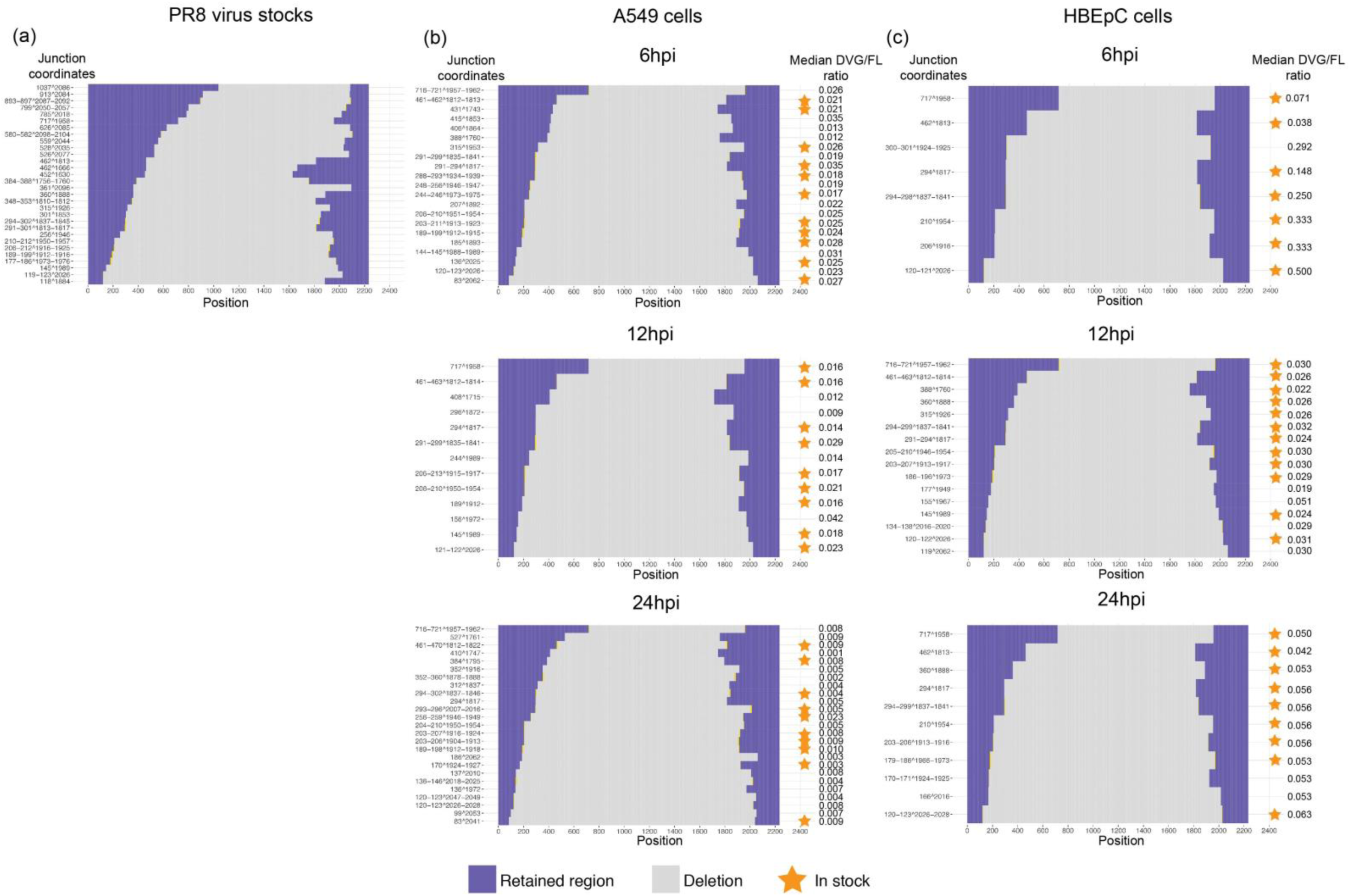
Distribution of junction sites in the PA segment and the medians of their DVG/FL ratio in cells where a given defective viral transcript was detected.

**Supplementary Figure 6.**
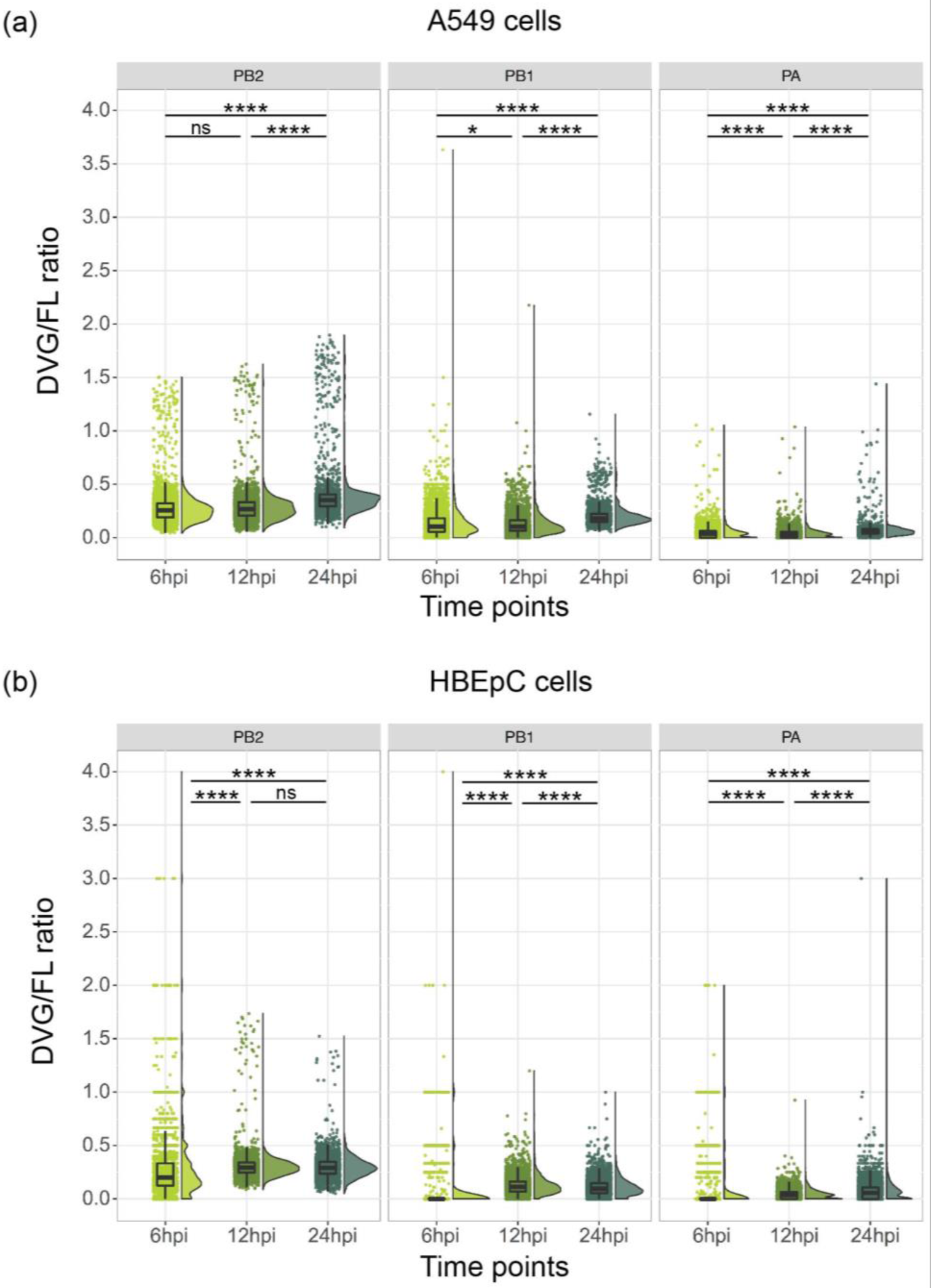
Distribution of the DVG/FL ratio in A549 and HBEPC cells over the course of the infection. All pairwise comparisons were performed with two-tailed Wilcoxon rank sum test, for which the null hypothesis was that cells harvested at two different time points have different median relative abundance of viral transcripts. The significance levels were denoted by the asterisks: * p ≤ 0.05, ** p ≤ 0.01, *** p ≤ 0.001, **** p ≤ 0.0001, and “ns” as not significant. All box plots show the first and third quantiles as the lower and upper hinges, the median in the center, and 1.5 * inter-quartile range (IQR) from the first and third quantiles as the whiskers.

**Supplementary Figure 7.**
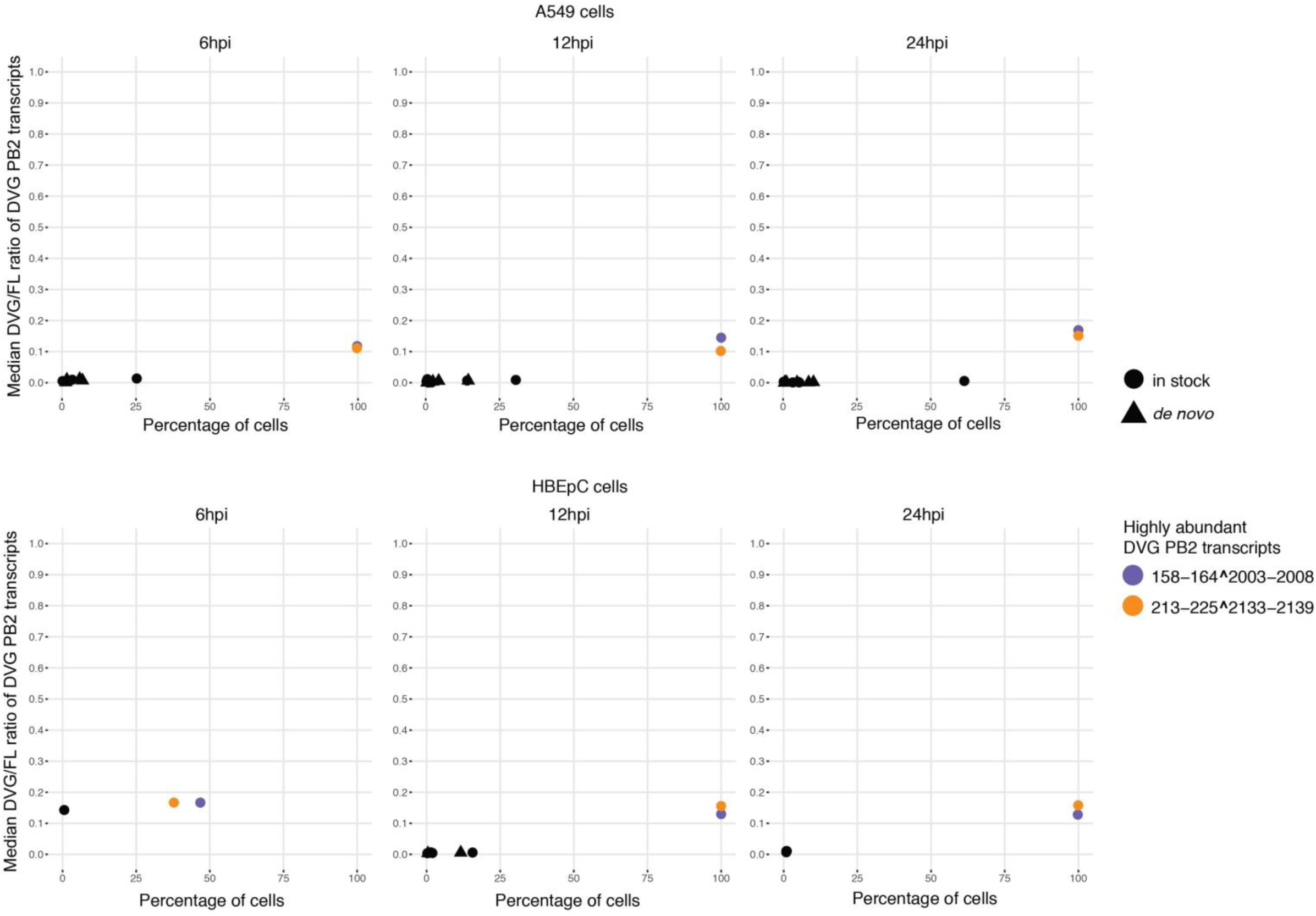
Percentage of A549 and HBEpC cells in which a given DVG PB2 transcript was detected versus the median of the DVG/FL ratio for that transcript in those cells over the course of the infection. Each type of DVG PB2 transcript was represented by a dot or triangle. A dot denoted the DVG transcripts also seen in the virus stock and a filled triangle denoted the *de novo* generated transcripts. Two dominant DVG PB2 species were highlighted in slate blue and orange.

**Supplementary Figure 8.**
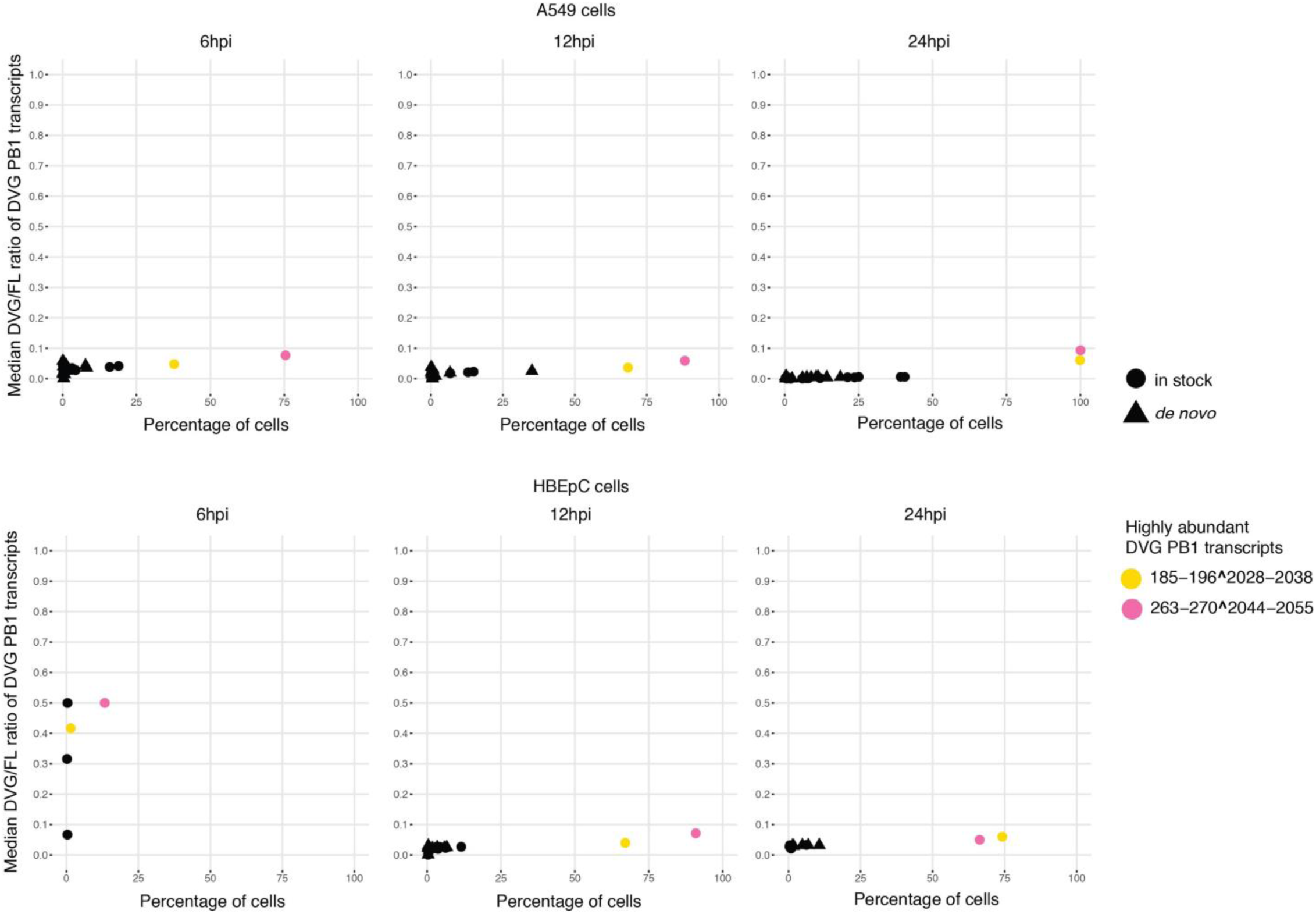
Percentage of A549 and HBEpC cells in which a given DVG PB1 transcript was detected versus the median of the DVG/FL ratio for that transcript in those cells over the course of the infection. Two dominant DVG PB1 species were highlighted in yellow and pink.

**Supplementary Figure 9.**
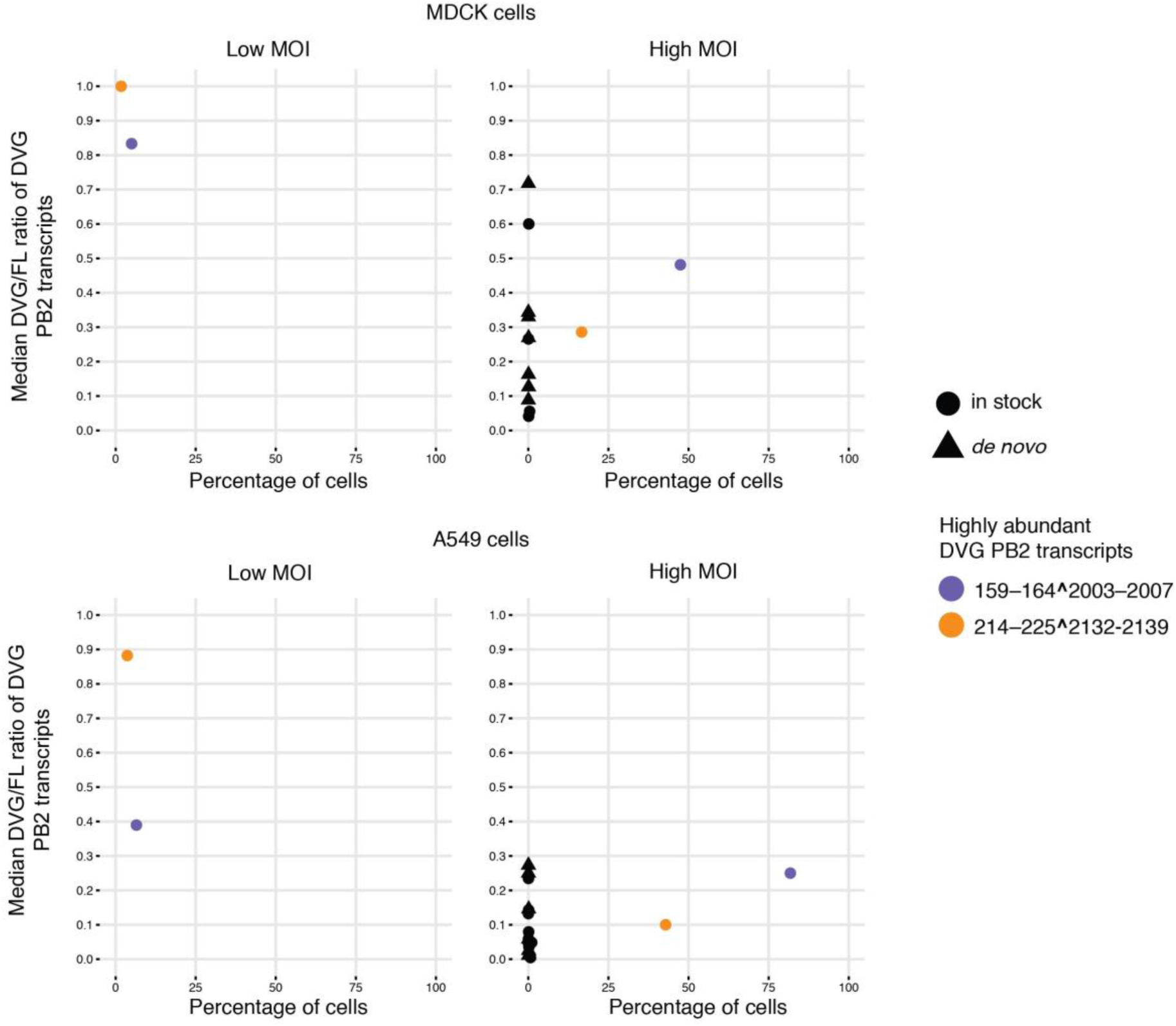
Percentage of MDCK and A549 cells in which a given DVG PB2 transcript was detected versus the median of the DVG/FL ratio for that transcript in those cells. MDCK and A549 cells were infected with the same PR8 virus stock at high (5) or low (0.2) MOI and harvested at 6hpi followed by 10X Genomics 3’ single-cell library preparation and sequencing. Two dominant DVG PB2 species were highlighted in slate blue and orange.

**Supplementary Figure 10.**
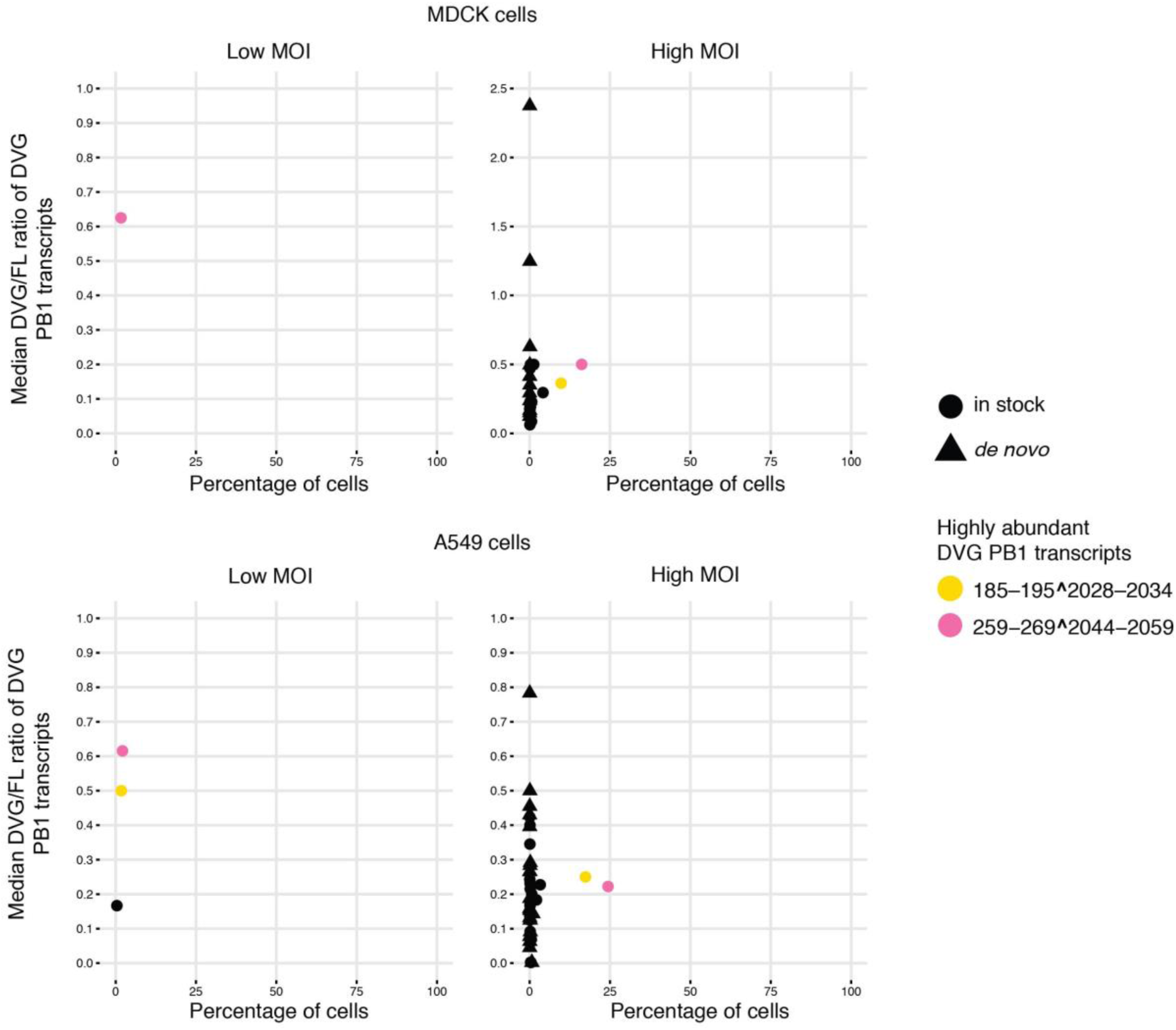
Percentage of MDCK and A549 cells in which a given DVG PB1 transcript was detected versus the median of the DVG/FL ratio for that transcript in those cells. MDCK and A549 cells were infected with the same PR8 virus stock at high (5) or low (0.2) MOI and harvested at 6hpi followed by 10X Genomics 3’ single-cell library preparation and sequencing. Two dominant DVG PB1 species were highlighted in yellow and pink.

**Supplementary Figure 11.**
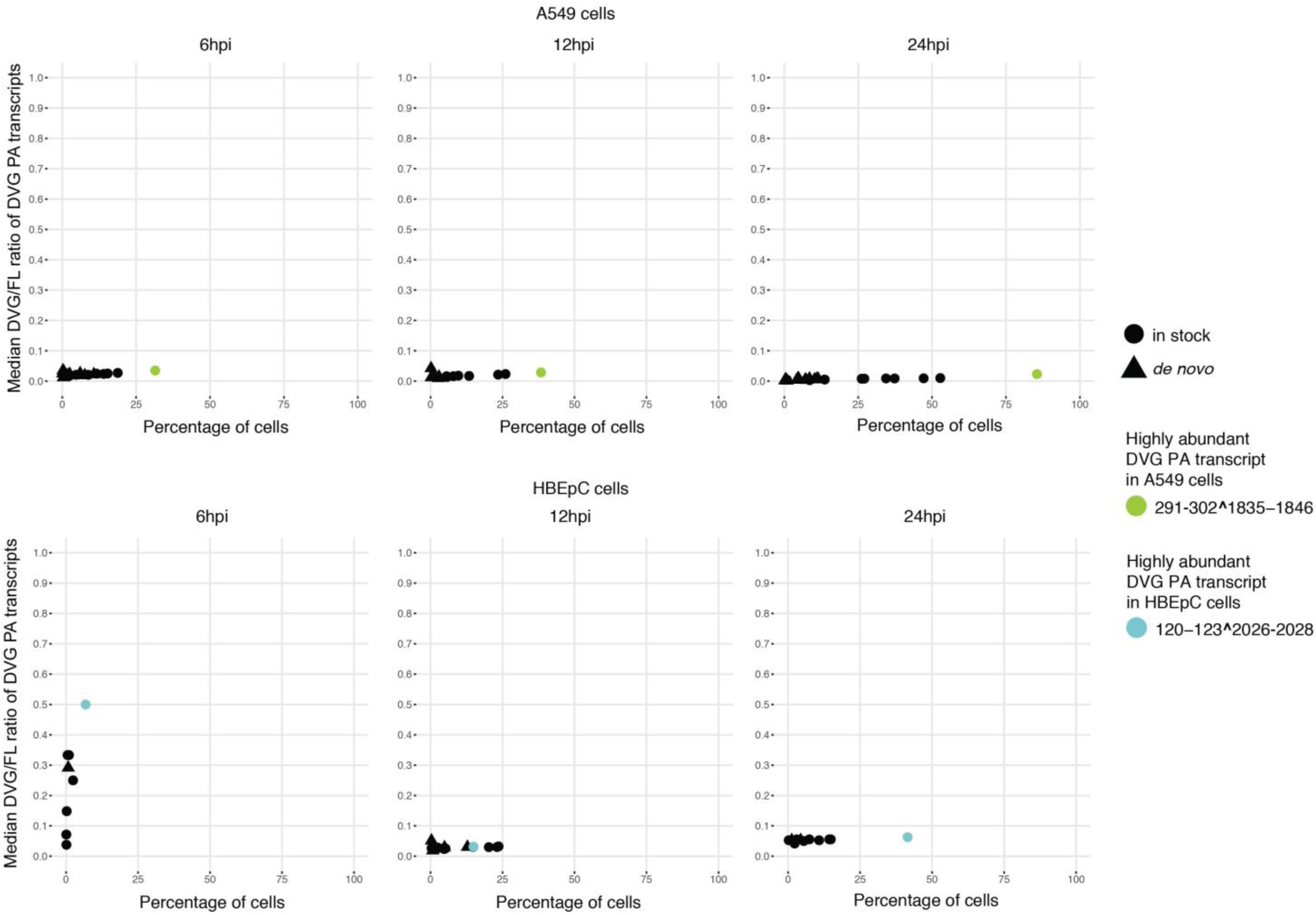
Percentage of A549 and HBEpC cells in which a given DVG PA transcript was detected versus the median of the DVG/FL ratio for that transcript in those cells over the course of the infection. The dominant DVG PA species in each cell type were highlighted.

**Supplementary Figure 12.**
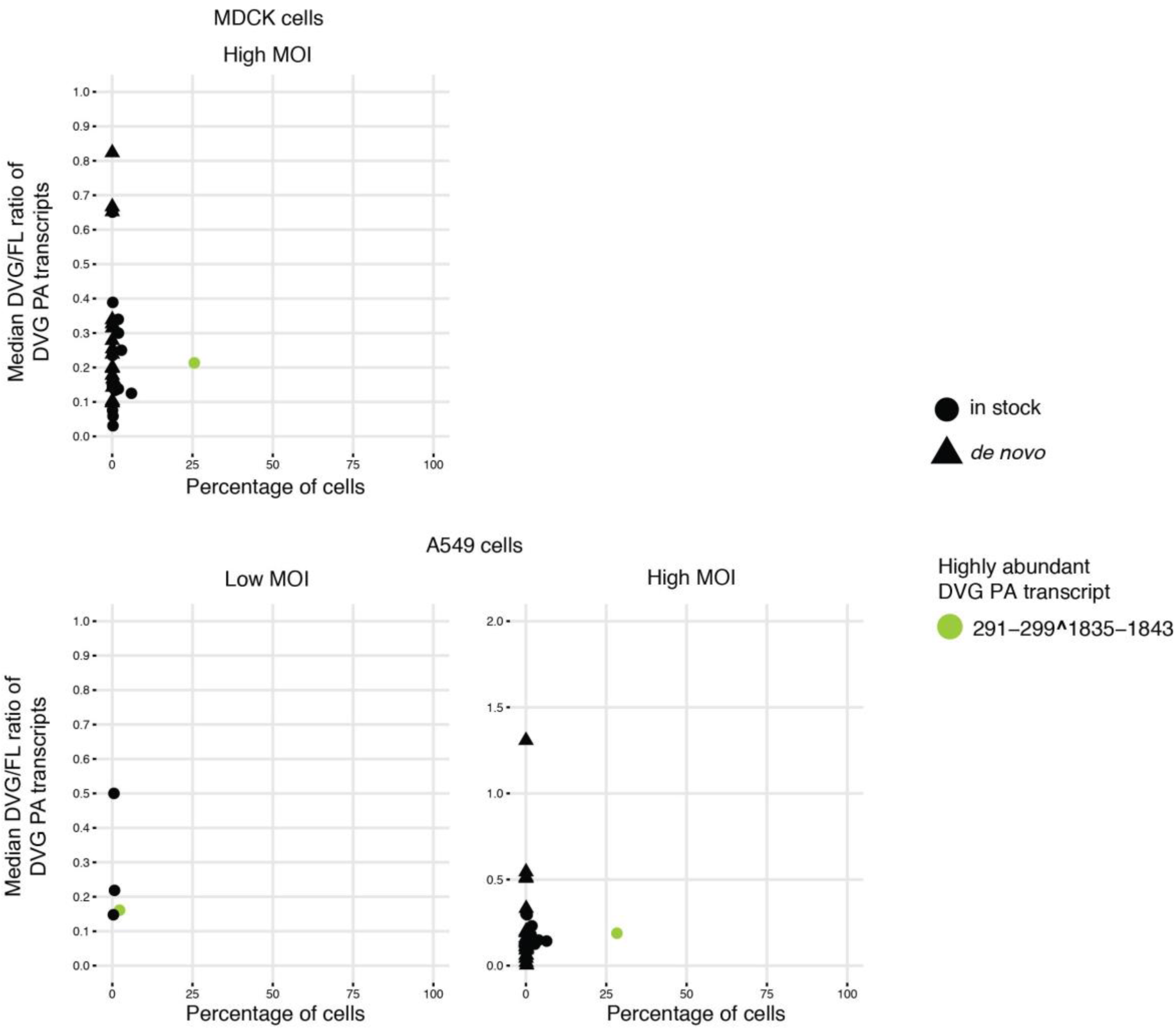
Percentage of MDCK and A549 cells in which a given DVG PA transcript was detected versus the median of the DVG/FL ratio for that transcript in those cells. MDCK and A549 cells were infected with the same PR8 virus stock at high (5) or low (0.2) MOI and harvested at 6hpi followed by 10X Genomics 3’ single-cell library preparation and sequencing. The DVG PA transcripts were not detected in MDCK cells infected at low MOI. The dominant DVG PA species was highlighted in chartreuse.

**Supplementary Figure 13.**
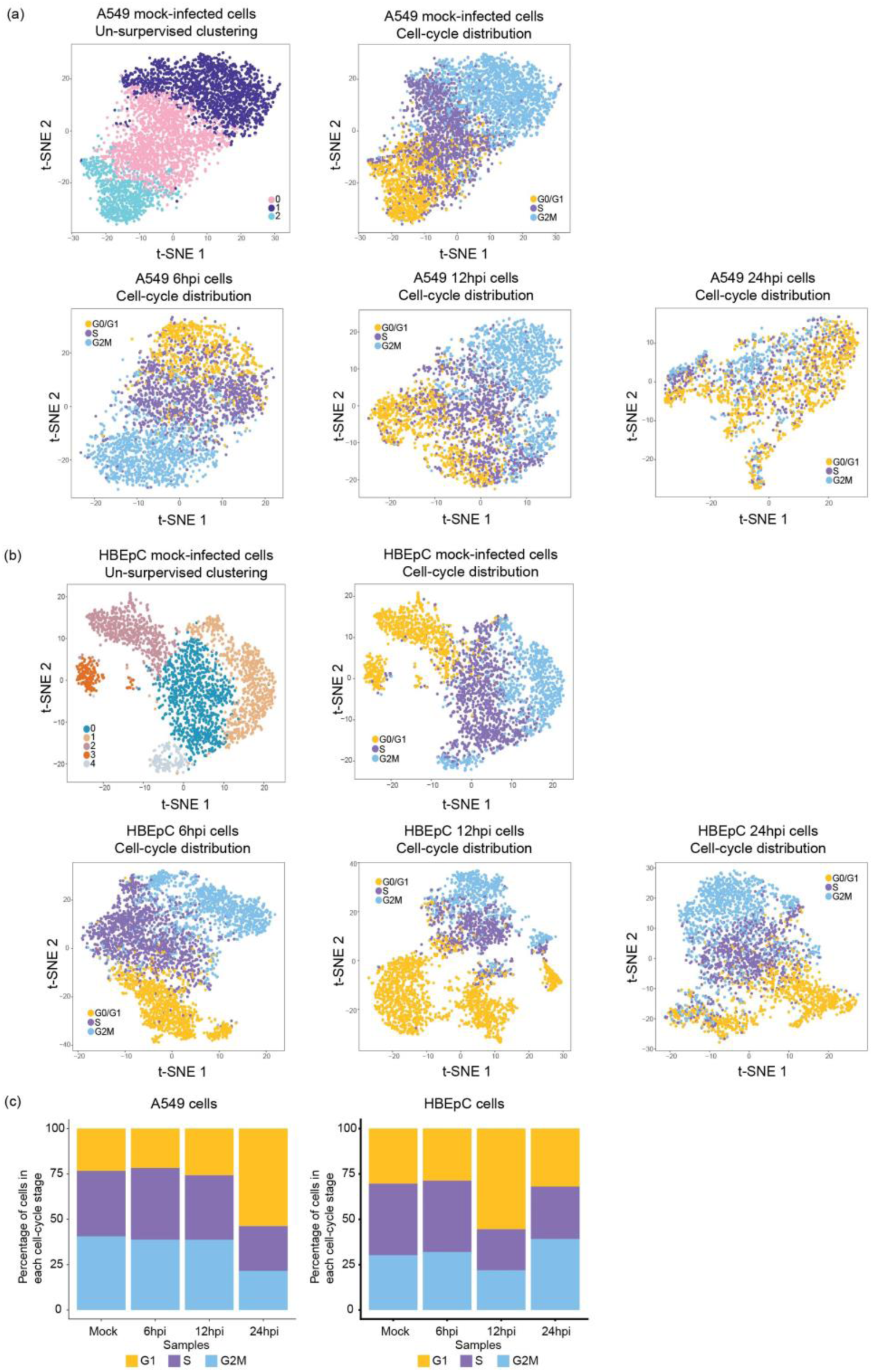
Visualization of un-supervised cell clustering on a t-SNE plot for mock-infected **(a)** A549 and **(b)** HBEpC cells and the distribution of cell-cycle within the population at each time point in two cell types. (**a**-**b**) Distribution of the cell-cycle stage for each cell represented by a dot on the t-SNE plot. Dots were colored by either the cluster identities for the mock-infected cells, or the cell-cycle stages for mock- and PR8-infected cells. The cell-cycle stage was assigned to each cell based on the expression level of a list of cell-cycle markers. (**c**) Distribution of the percentage of cells in each cell-cycle stage in two cell types.

**Supplementary Figure 14.**
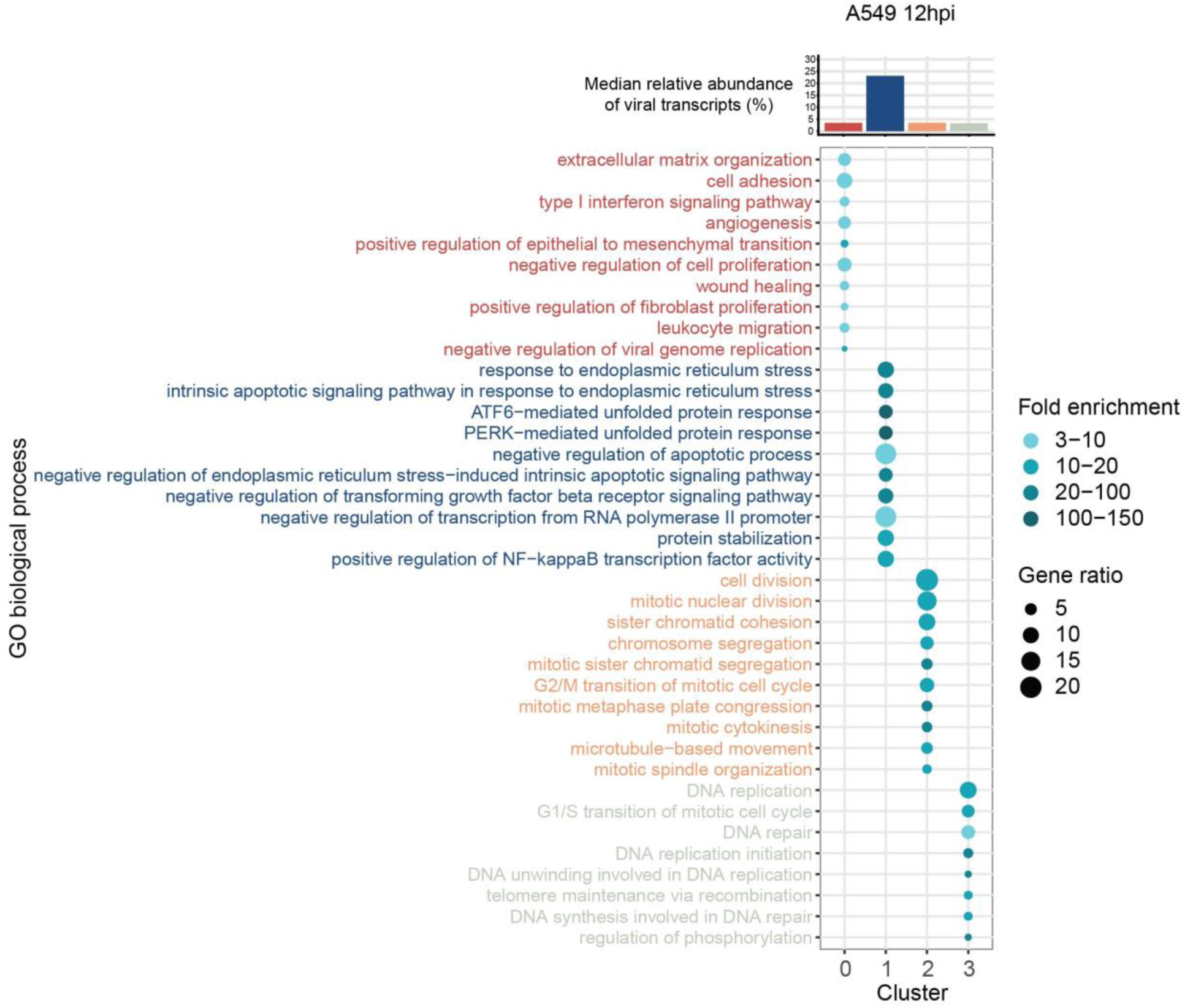
Enriched biological process gene ontology (GO) terms in A549 at 12hpi. The top 10 over-represented GO terms in each cluster are denoted by dots. The color intensity of each dot indicates the fold enrichment of the corresponding GO term and the size corresponds to the ratio of queried genes in the gene set associated with a given GO term. The GO terms enriched in each cluster are color-coded by clusters. The bar chart above the bubble chart of the GO terms shows the median relative abundance of the viral transcripts across cells in each cluster. The colors of the bar are consistent with that of the GO terms.

**Supplementary Figure 15.**
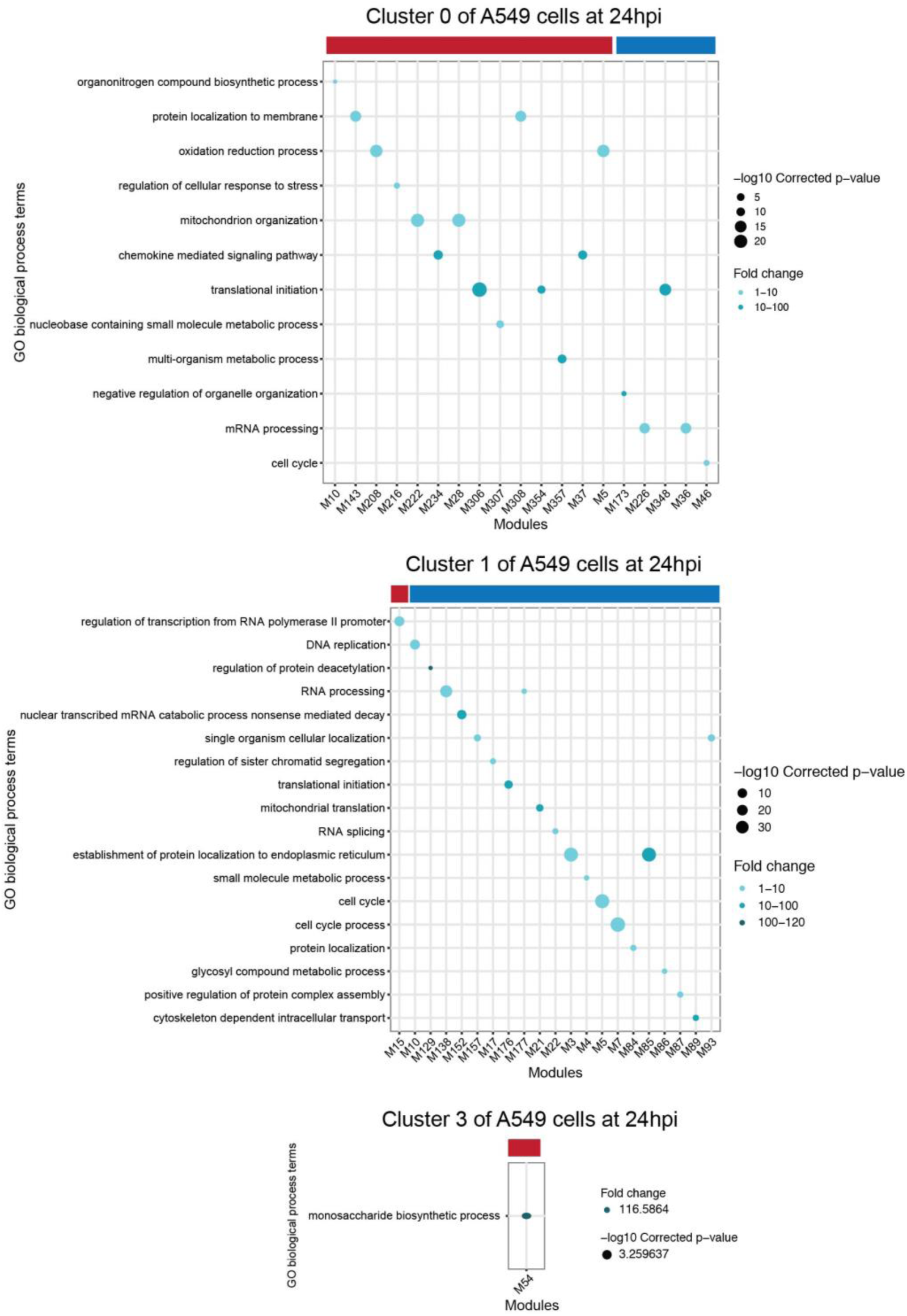
Functional MEGENA modules in clusters 0 and 1 of A549 at 24hpi. Each panel shows enriched biological process GO terms in the modules correlated with the relative abundance of virus transcripts in corresponding clusters. Each GO term is denoted by a dot. The color intensity of each dot indicates the fold enrichment of the corresponding GO term and the size corresponds to the -log10 transformed corrected p-value for a given GO term. Red and blue color-bars above the dot plot denote positive or negative correlation of viral transcription with corresponding modules, respectively. Modules that have a significant correlation with the relative abundance of virus transcripts and enriched GO biological process terms are shown in the GO term plots. Module member information for all viral transcription correlated-modules can be found on Github (https://github.com/GhedinLab/Single-Cell-IAV-infection-in-monolayer/tree/master/SupplementaryTables).

**Supplementary Figure 16.**
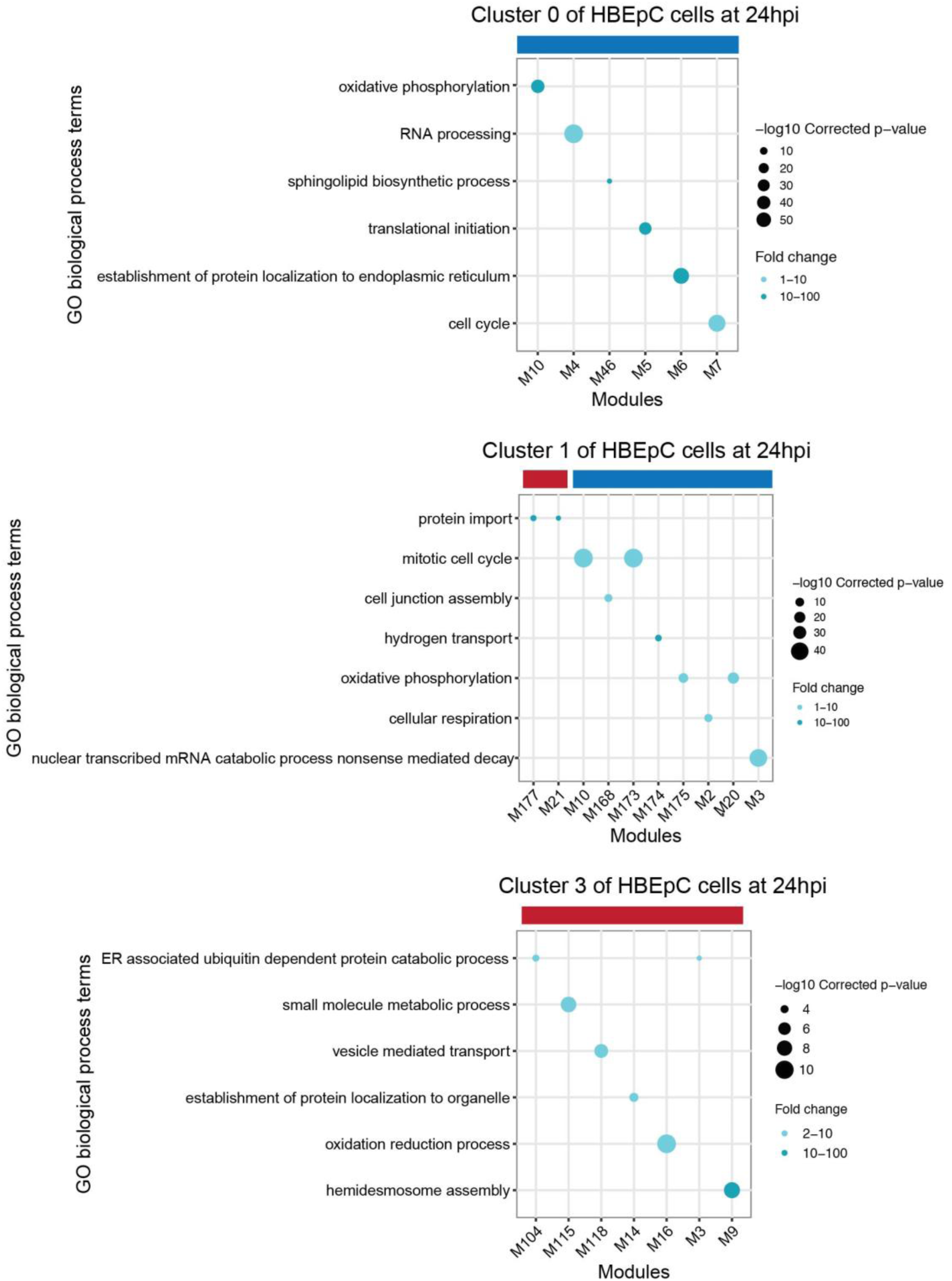
Functional MEGENA modules in clusters 0, 1, and 3 of HBEpC at 24hpi. Each panel shows enriched biological process GO terms in the modules correlated with the relative abundance of virus transcripts in corresponding clusters. Each GO term is denoted by a dot. The color intensity of each dot indicates the fold enrichment of the corresponding GO term and the size corresponds to the -log10 transformed corrected p-value for a given GO term. Red and blue color-bars above the dot plot denote positive or negative correlation of viral transcription with corresponding modules, respectively. Modules that have a significant correlation with the relative abundance of virus transcripts and enriched GO biological process terms are shown in the GO term plots. Module member information for all of viral transcription correlated-modules can be found on Github (https://github.com/GhedinLab/Single-Cell-IAV-infection-in-monolayer/tree/master/SupplementaryTables).

**Supplementary Figure 17.**
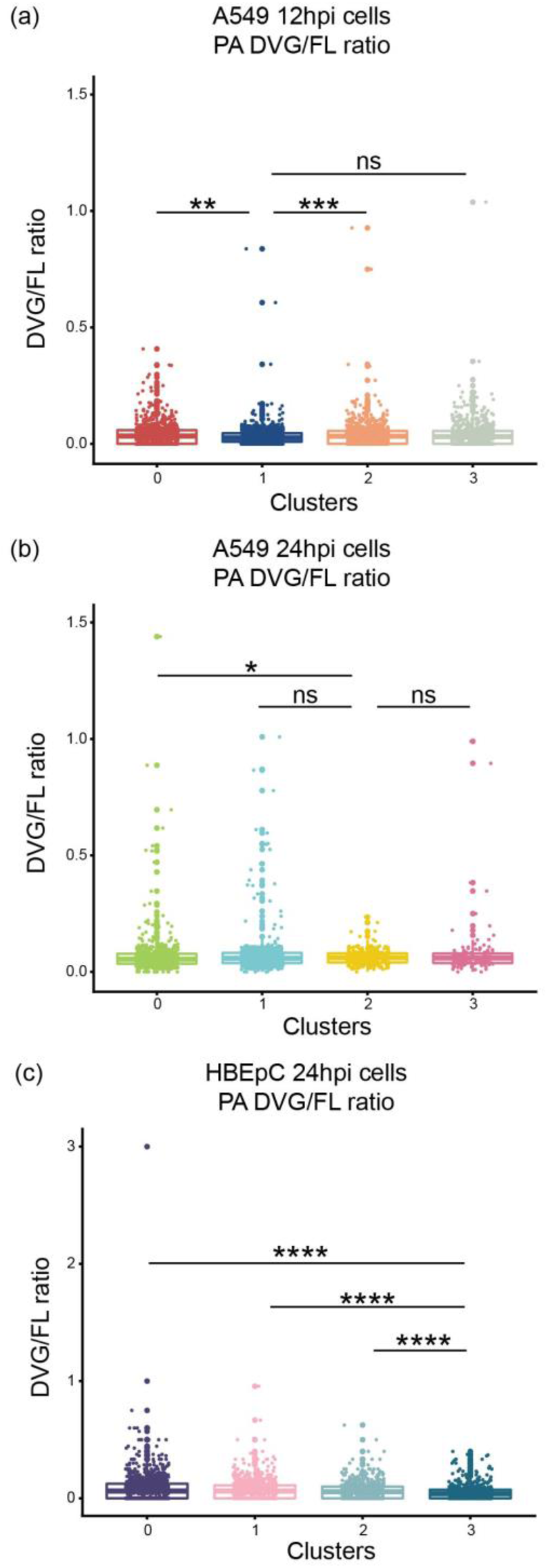
Distribution of the DVG/FL ratio for the DVG PA transcripts in each cluster of A549 cells at 12hpi and 24hpi and HBEpC cells at 24hpi. All box plots show the first and third quantiles as the lower and upper hinges, the median in the center, and 1.5 * inter-quartile range (IQR) from the first and third quantiles as the whiskers. The significance levels of pairwise comparisons determined by one-tailed Wilcoxon rank sum test were denoted by the asterisks: * p ≤ 0.05, ** p ≤ 0.01, *** p ≤ 0.001, **** p ≤ 0.0001, and “ns” as not significant.

**Supplementary Figure 18.**
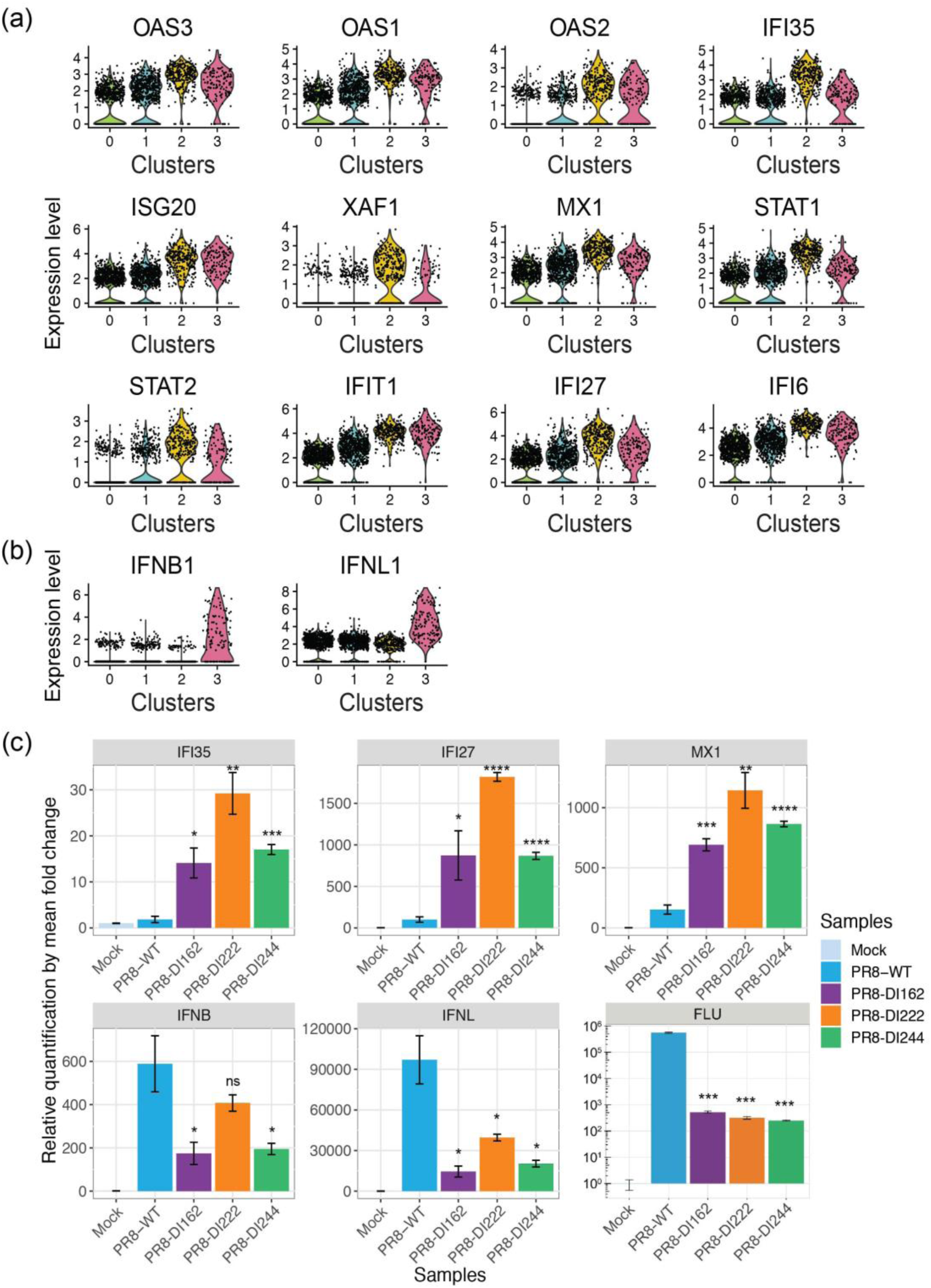
Expression of ISGs and IFNs in DI-enriched A549 cells at 24hpi. **(a-b)** The expression level of **(a)** significantly over-expressed genes in cluster 2, which are associated with GO terms related to the type I IFN signaling pathway and have the fold change on the log-scale >= 1, and **(b)** type I and III IFNs, in each cluster of A549 cells at 24hpi. The dot represents the normalized expression level of a gene in each cell. The violin shade is colored by the cluster identity. **(c)** Validation of over-expression of selected ISGs, IFNs, and the M gene of IAV (i.e., FLU) by qPCR. Subconfluent A549 cells were either mock-infected or infected with one type of PR8-DI virus (PR8-DI162, -DI222, or -DI244) or PR8-WT virus at a MOI of 10. Total RNA was extracted from cells collected at 24hpi. The levels of ISGs, IFNs, and the IAV M gene mRNA were determined by qPCR and normalized to the level of β-actin (ACTB). The error bar representing the standard deviation of the mean was obtained from three biological replicates. Statistical tests were done using two-tailed Student’s *t*-test in R to compare the differences in fold change between PR8-WT and PR8-DI infection. The significance levels were denoted by the asterisks: * p ≤ 0.05, ** p ≤ 0.01, *** p ≤ 0.001, **** p ≤ 0.0001, and “ns” as not significant.

**Supplementary Figure 19.**
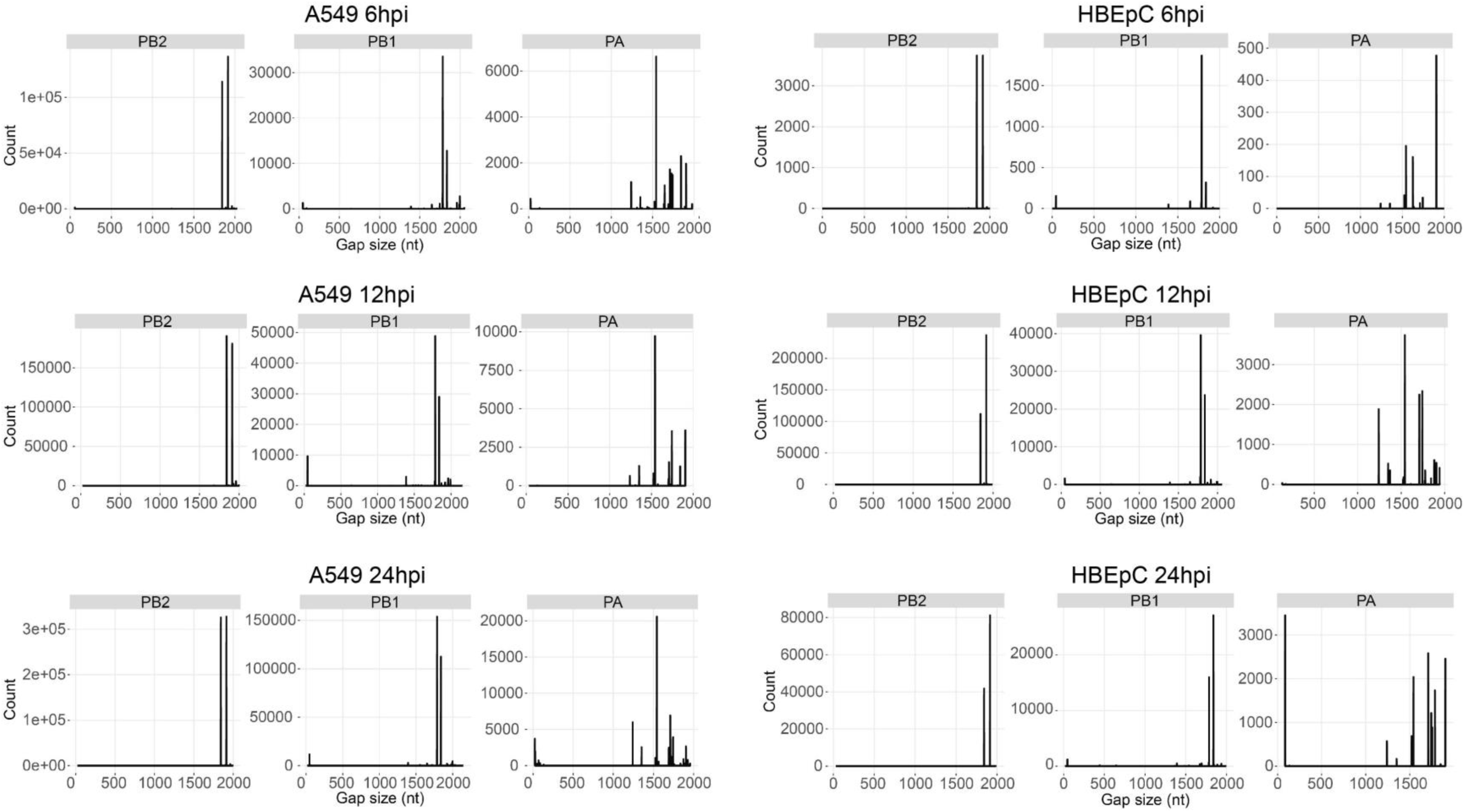
Distribution of the size of the deletions in the gap-spanning reads aligned to the polymerase segments in two cell types over the course of the infection. The frequency of each size detected in a sample was represented by the height of the bar shown on the y-axis.

